# Injectable MSC Spheroid and Microgel Granular Composites for Engineering Cartilage Tissue

**DOI:** 10.1101/2022.12.28.522141

**Authors:** Nikolas Di Caprio, Matthew D. Davidson, Andrew C. Daly, Jason A. Burdick

## Abstract

Articular cartilage is important for joint function, yet it can be damaged due to disease or trauma. Cartilage lacks natural healing processes and current clinical treatments are limited in their ability to form functional cartilage for repair. Here, we reimagine cartilage tissue engineering with an approach that combines aggregates of adult MSCs (spheroids) with hydrogel microparticles (microgels) to form granular composites that are injectable, permit cell-cell contacts for chondrogenesis, allow spheroid fusion and growth, and undergo interparticle crosslinking post-injection via light for stability. We use simulations and experimental analyses to establish the importance of initial MSC spheroid to microgel volume ratios in granular composites that balance mechanical support with tissue growth. Long-term chondrogenic cultures of granular composites produce engineered cartilage tissue within the range of native properties, which can be further enhanced via MSC/chondrocyte co-cultures. Altogether, we have developed a new strategy of injectable granular composites for engineering cartilage tissue.

---

Damage to articular cartilage and progression to disease is a prevalent problem that can limit joint mobility and result in joint pain for patients, reducing quality of life.^1^ Cartilage damage in the context of focal lesions can develop from physical trauma and when left untreated result in joint inflammation and matrix degradation, initiating the onset of osteoarthritis (OA) (Fig. 1a).^2^ Cartilage tissue lacks the innate ability to repair itself due to its avascular nature and requires clinical strategies for repair when damaged. Such strategies that use chondrocytes (CHs) for repair (e.g., matrix-induced/autologous chondrocyte implantation) suffer limitations of cell sourcing from healthy cartilage, dedifferentiation of chondrocytes during expansion, and mechanically inferior cartilage repair tissue.^3,4^ These drawbacks motivate the use of mesenchymal stromal cells (MSCs), isolated primarily from bone marrow or adipose tissue, as an alternative cell source to repair damaged cartilage.

**Fig. 1.**
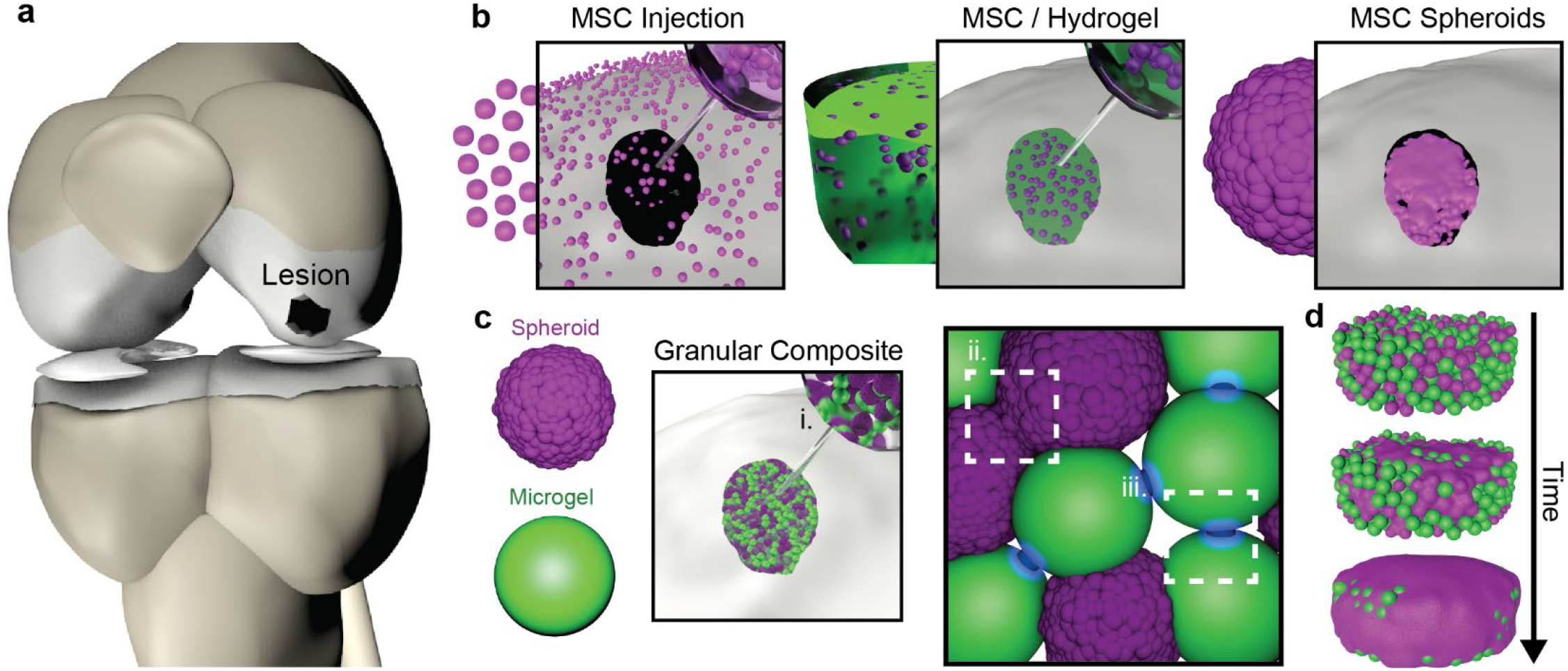
Granular composites address limitations of current MSC-based cartilage repair strategies. **a**, Schematic representation of an articular cartilage focal defect. **b**, Current MSC-based cartilage repair strategies: injection of MSC suspensions, which suffer from low cell retention within the defect; the encapsulation of MSCs within hydrogels (MSC/Hydrogel), which improve cell retention, but limit cell-cell contacts and ECM distribution; and MSC spheroids, which permit cell-cell contacts, but lack mechanical support within the defect and can result in collapse. **c**, Schematic overview of granular composite design, where MSC spheroids and NorHA microgels are mixed to enable (i) injectability for delivery to defects, (ii) cell-cell contacts for enhanced chondrogenesis and spheroid fusion for tissue formation, and (iii) interparticle crosslinking via light to stabilize the composite when desired. **d**, Schematic of granular composites over time, where spheroid fusion and growth result in cartilage tissue throughout the granular hydrogel.

One approach for MSC-based cartilage repair includes intra-articular injection of MSCs in suspension; however, insufficient cell retention within the defect and fibrocartilage production limit therapeutic efficacy, and despite numerous clinical trials, no Food and Drug Administration approved products are available with this approach (Fig. 1b).^5–9^ To improve these outcomes, MSCs are embedded within hydrogels to enhance their localization to cartilage defects and to provide an environment that supports MSC chondrogenesis; however, many hydrogels limit the distribution of deposited extracellular matrix (ECM) and restrict important cell-cell contacts that are known to enhance MSC chondrogenesis and improve cartilage formation.^10–15^ In contrast, aggregates of MSCs (i.e., spheroids) enable cell-cell contacts and high-cell density cultures that enhance cartilaginous ECM deposition;^16^ yet, individual spheroids are insufficient to fill defects and spheroid suspensions lack the mechanical support to limit fusion into a collapsed pellet.^17,18^ Spheroids from CHs have been investigated via implantation into defects, but suffer from limitations of long pre-culture periods and collapse within lesions.^19^ Ultimately, spheroids provide positive cell-cell contacts to enhance cartilage formation, but lack mechanical support; whereas, hydrogels introduce structural support, but limit cell-cell contacts and ECM distribution to enable functional tissue properties.

Granular hydrogels are an emerging class of hydrogels that consist of packed hydrogel microparticles (i.e., microgels), resulting in properties such as injectability due to microgel flow, intrinsic porosity via microgel packing, and mechanical support via interparticle crosslinking. Such hydrogels permit cellular migration within the pore structure, particularly compared to traditional uniform hydrogels.^20–22^ Here, we combine the positive attributes of MSC spheroids and granular hydrogels to engineer cartilage through the formation of granular composites. Specifically, adult MSC spheroids are mixed with norbornene-modified hyaluronic acid (NorHA) microgels to form granular composites that (i) are injectable for delivery to molds or defects, (ii) introduce cell-cell contacts and spheroid fusion that are conducive to cartilage formation, and (iii) undergo interparticle crosslinking via light to immediately stabilize constructs (Fig. 1c). Over time, MSC chondrogenesis and ECM deposition facilitate the formation of cartilage tissue that is guided by the stabilized granular hydrogel (Fig. 1d). Further, we introduce MSC and CH co-culture spheroids into our granular composites to enhance MSC chondrogenesis and cartilage formation, motivated by prior studies that show the benefits of such co-culture platforms.^23,24^

## Results

### Spheroids and microgels form granular composites when mixed

Before investigating granular composite cartilage formation, we first sought to understand how individual MSC spheroids grow in chondrogenic medium. MSC spheroids are fabricated by seeding MSCs into pyramidal microwells that allow cells to condense by day 2 (D2) (Supplemental Fig. 1a). These microwells allow for scalable generation of spheroids in the thousands, where the initial MSC seeding density (500-2000 cells/spheroid) controls the MSC spheroid mean diameter (∼125-175 µm) and distribution (Supplemental Fig. 1b). Spheroid viability is high, with a live cell area >90% after D7, except for the largest MSC spheroids (2000 cells/spheroid) with a live cell area of ∼80% (Supplemental Fig. 2). To incorporate the highest cell density while also maintaining high cell viability, we select the 1000 cell/spheroid seeding density for the remaining studies. MSC spheroids increase ∼25% in diameter and ∼100% in volume (assuming a perfect sphere) over 28 days (Supplemental Fig. 3a, b). Staining for cartilage ECM markers chondroitin sulfate and collagen II reveal increased ECM deposition over time, indicating MSC spheroid chondrogenesis coinciding with increased spheroid size (Fig. 2b). ECM deposition is further confirmed via histological staining for collagens and sulfated glycosaminoglycans (sGAGs) (Supplemental Fig. 3c).

**Fig. 2.**
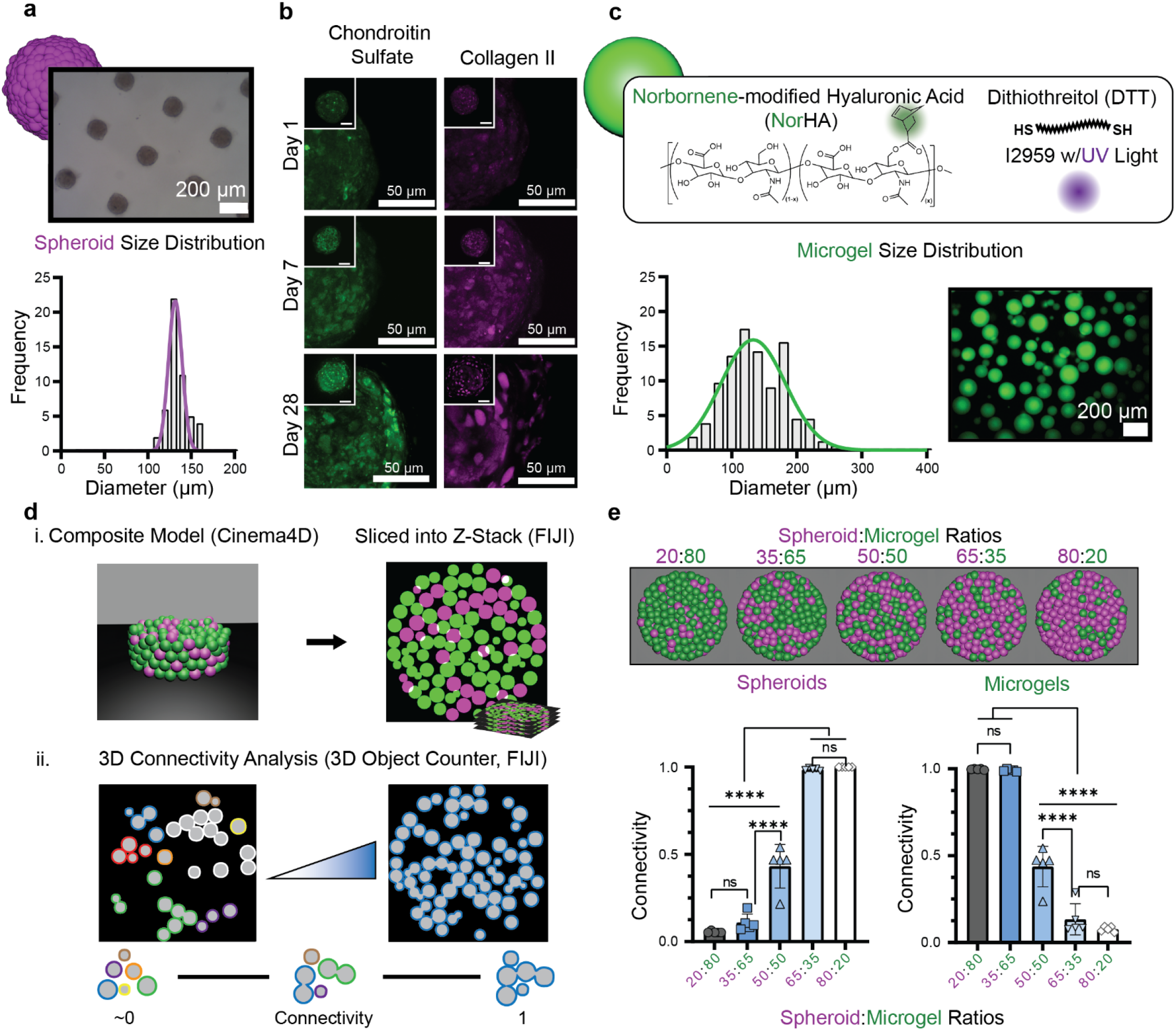
Spheroids and microgels form granular composites and their ratio dictates connectivity. **a**, Representative image of MSC spheroids (top) and quantification of MSC spheroid size distribution (bottom) 2 days after seeding 1000 cells/spheroid. n=50 spheroids combining 3 biologically independent donors; scale bar: 200μm. **b**, Representative images of MSC spheroid immunostaining for collagen II (magenta) and chondroitin sulfate (green) over 28 days; scale bars: 50µm. **c**, Quantification of microgel size distribution (left) and representative image (right) of microgels fabricated via batch emulsion of NorHA (spun at 280 RPM). n=154 from 2 independent batch emulsions; scale bar: 200μm. **d**, Schematic workflow of dynamic rigid body simulation connectivity analysis through Cinema4D. (i) Granular composite mixing (green: microgels; pink: spheroids) simulated via gravity and imported into FIJI to slice and voxelize 3D object distributions as binary Z-stacks. (ii) Binary Z-stacks analyzed for connectivity through 3D object counter function in FIJI. Connectivity (ranges: approaching 0 to 1) of an object type (spheroid or microgel) is defined as the average number of objects within “clusters” (represented as distinct colors and defined as continuous touching of objects in 3D) divided by the total number of individual objects within the composite (1: all objects connected in 3D, approaching 0: no objects connected in 3D). **e**, Representative images (top) and quantified connectivity (bottom; left: spheroids, right: microgels) of granular composite simulations of varying volume ratios of spheroids to microgels. n=5 independent simulations, mean ± s.d.

After characterizing individual MSC spheroids, we next fabricate similar-sized microgels. Due to the presence of hyaluronic acid (HA) within cartilage and the long history of HA in biomedical products (e.g., viscosupplements) and in hydrogel design, we use HA with norbornene modification (i.e., NorHA) to form microgels.^25–27^ We fabricate NorHA microgels using batch emulsion, by adding a solution of NorHA, crosslinker, and photoinitiator dropwise into mineral oil while mixing. The emulsion is exposed to UV light for 30 min. for microgel crosslinking and microgel suspensions are obtained by washing from oil (Supplemental Fig. 1c). Rotation spin speeds alter the mean diameters and distributions of NorHA microgels (e.g., increased speeds reduce NorHA microgel size) (Supplemental Fig. 1d). To match MSC spheroid diameters, an emulsion spin speed of 280 RPM is used to obtain NorHA microgel diameters of ∼140 µm (Fig. 2c).

With each granular component characterized, we next investigate how the mixing of spheroids and microgels influences granular composite formation and the connectivity of each component. This is important to support both spheroid fusion and growth for tissue formation and microgel interparticle crosslinking for stabilization. To gain insight into this and to avoid extensive iterative experimental testing, mixing is simulated in Cinema4D (C4D).^28,29^ We hypothesize that microgels within the composite require a critical level of connectivity to maintain construct stability, below which the composite will disassemble upon agitation. The mixing of spheroids and microgels within a cylindrical tube is simulated in C4D, where gravity drives mixing behaviors (Supplemental Video 1). Voxelation of simulated composites allows for generation of a binary z-stack that is then analyzed in FIJI and the two components are separated (Fig. 2di). The 3D object counter then assigns colors to “clusters” of either spheroids or microgels that are physically touching in 3D space, allowing the calculation of a connectivity parameter for either granular component (defined as average number of spheroids or microgels in “clusters” divided by the total number of spheroids or microgels within the composite) (Fig. 2dii) (Supplemental Video 2). If all spheroids or microgels are connected, the connectivity parameter would be 1, whereas if there are low numbers in each cluster, the connectivity parameter approaches 0. Similar to percolation theory, this method allows us to understand the connectivity of both spheroid or microgel components to inform how the initial volume ratio impacts stability and tissue formation.^30^ This approach reveals that low ratios (<20:80) of spheroids to microgels result in primarily isolated spheroids (i.e., that would not result in fusion) with low spheroid connectivity, whereas high ratios of spheroids to microgels (>50:50) result in decreased microgel contact (i.e., limited interparticle crosslinking) with low microgel connectivity (Fig. 2e). Further, microgel size polydispersity and spheroid aggregation are introduced into simulations and demonstrate reduced microgel connectivity with polydisperse microgels and enhanced spheroid connectivity with aggregation (Supplemental Fig. 4). Taken together, we identify 20:80, 35:65, and 50:50 spheroid to microgel volume ratio formulations for further exploration of granular composites, based on the need for high microgel connectivity for construct stabilization and to limit issues of construct collapse over time with spheroids alone.^18,31^

### Granular hydrogels are injectable and mechanically stabilized with interparticle crosslinking

To investigate potential injectability of granular composites, the rheological properties of 20:80, 35:65, and 50:50 spheroid to microgel ratios are measured. Spheroids and microgels are first independently isolated and concentrated via centrifugation and then homogenously mixed to form a composite slurry (Fig. 3a). As controls, either MSC spheroids alone or microgels alone are measured rheologically (Fig. 3b), where the microgels exhibit a 10-fold increase in storage modulus and an increase in strain yielding properties when compared to spheroids (Fig. 3b). When composites are tested, the composition reflects the properties with reduced moduli and strain yielding behavior with increased amounts of spheroids (Fig. 3c). The quantified values for the storage moduli across groups are similar when microgels are included (except 20:80), whereas the loss modulus increases in composites over either component alone (Fig. 3d, e). It should be noted that the jamming conditions (e.g., centrifugation speed) of microgels can alter microgel packing and storage and loss moduli and must be considered when formulating composites and as expected, exposure to light stabilizes jammed microgels (Supplemental Fig. 5a, b). Cyclic strain behavior of individual components alone and granular composites across all volume ratios demonstrate shear-thinning and self-healing properties (Supplemental Fig. 5c) and representative composites are easily ejected from a syringe and injected into a model cartilage defect (Supplemental Videos 3,4). Overall, these results indicate that all granular composites and their individual components are injectable and that composite properties are dependent on the formulation, likely due to the differences in properties between the elastic NorHA microgels and the viscoelastic spheroids.^32^

**Fig. 3.**
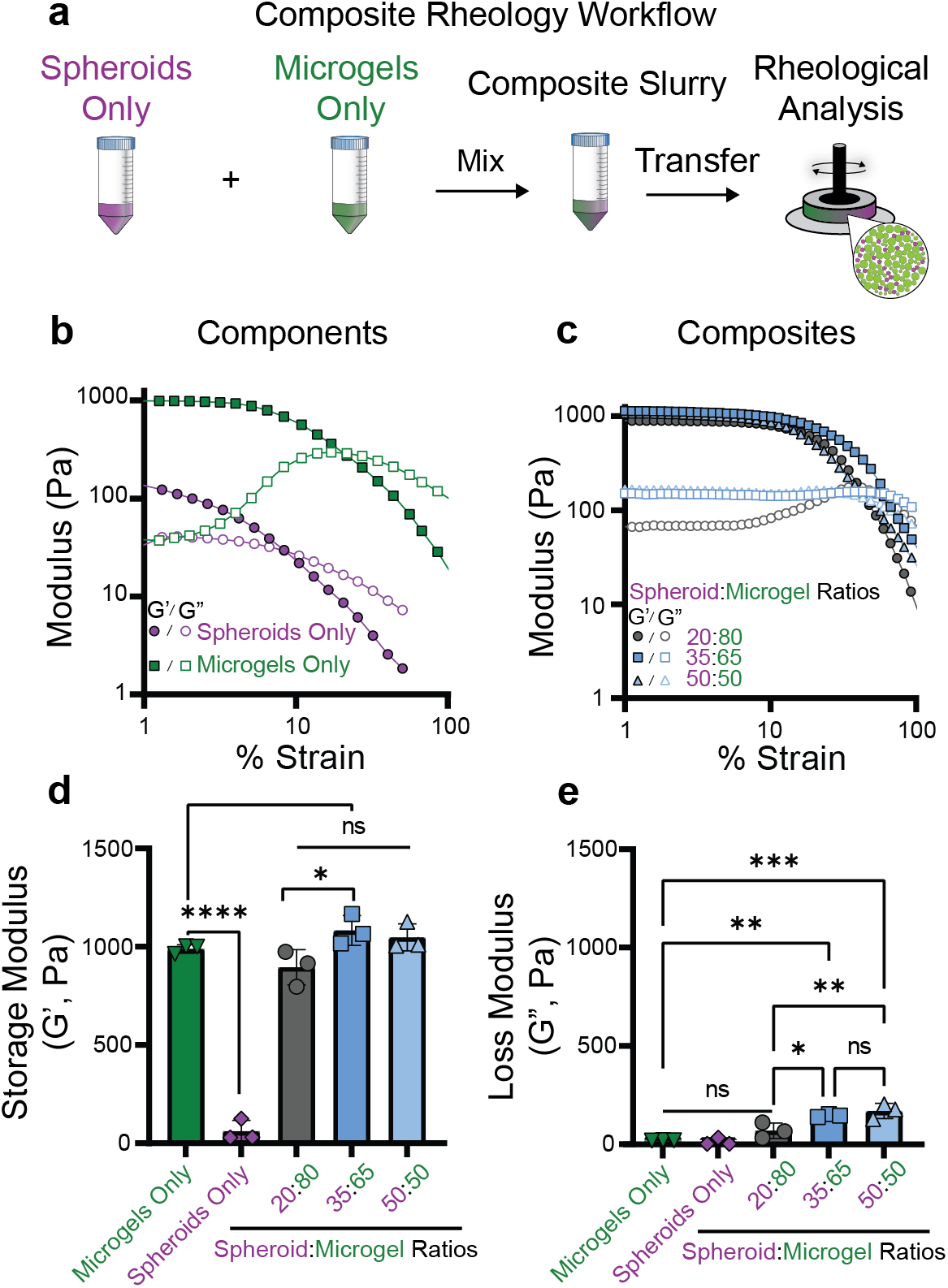
Granular hydrogels are injectable with properties based on formulation. **a**, Schematic of the composite rheology workflow. MSC spheroids and NorHA microgels are jammed independently, mixed into a composite slurry, and transferred onto a rheometer for analysis. **b**, Representative strain sweeps (1 Hz) of spheroid only and microgel only granular components. **c**, Representative strain sweeps (1 Hz) of granular composites at varying volume ratios of spheroids to microgels. Filled objects: G’, Empty objects: G”. **d**, Quantification of average storage moduli of granular components or granular composites with varying volume ratios. n= 3 independent rheological runs, mean ± s.d. **e**, Quantification of average loss moduli of granular components or granular composites with varying volume ratios. n= 3 independent rheological runs, mean ± s.d.

Based on the granular composite simulation findings and the injectability across these formulations, granular composites with 20:80, 35:65 and 50:50 spheroid to microgel ratios are fabricated by transferring composite slurries into a 3D printed mold, exposing to visible light for 3 min. for microgel interparticle crosslinking, and removing from molds for culturing (Fig. 4a). To visualize granular composites, constructs are cleared with RapiClear® to visualize whole MSC spheroids, confocal z-stacks are captured, and 3D reconstructions are performed on Imaris microscopy software (Fig. 4b). Upon hydrating composites, the 50:50 granular composite lacks mechanical support, likely due to low microgel connectivity that is insufficient to form a stable construct (Supplemental Fig. 6). Despite this, both 20:80 and 35:65 granular composites are mechanically stable after hydration. Expected volume ratios are validated from 3D confocal reconstructions of both groups, resulting in volume ratios within 3-5% on average of anticipated values within granular composites (Fig. 4c). Both 20:80 and 35:65 groups maintain a porosity volume of ∼20% (Fig. 4d). The pore area (750-1250 µm^2^) is significantly lower in the 35:65 group compared to the 20:80 group, likely due to the increased spheroid concentration and corresponding deformability of the composite during fabrication (Fig. 4d).

**Fig. 4.**
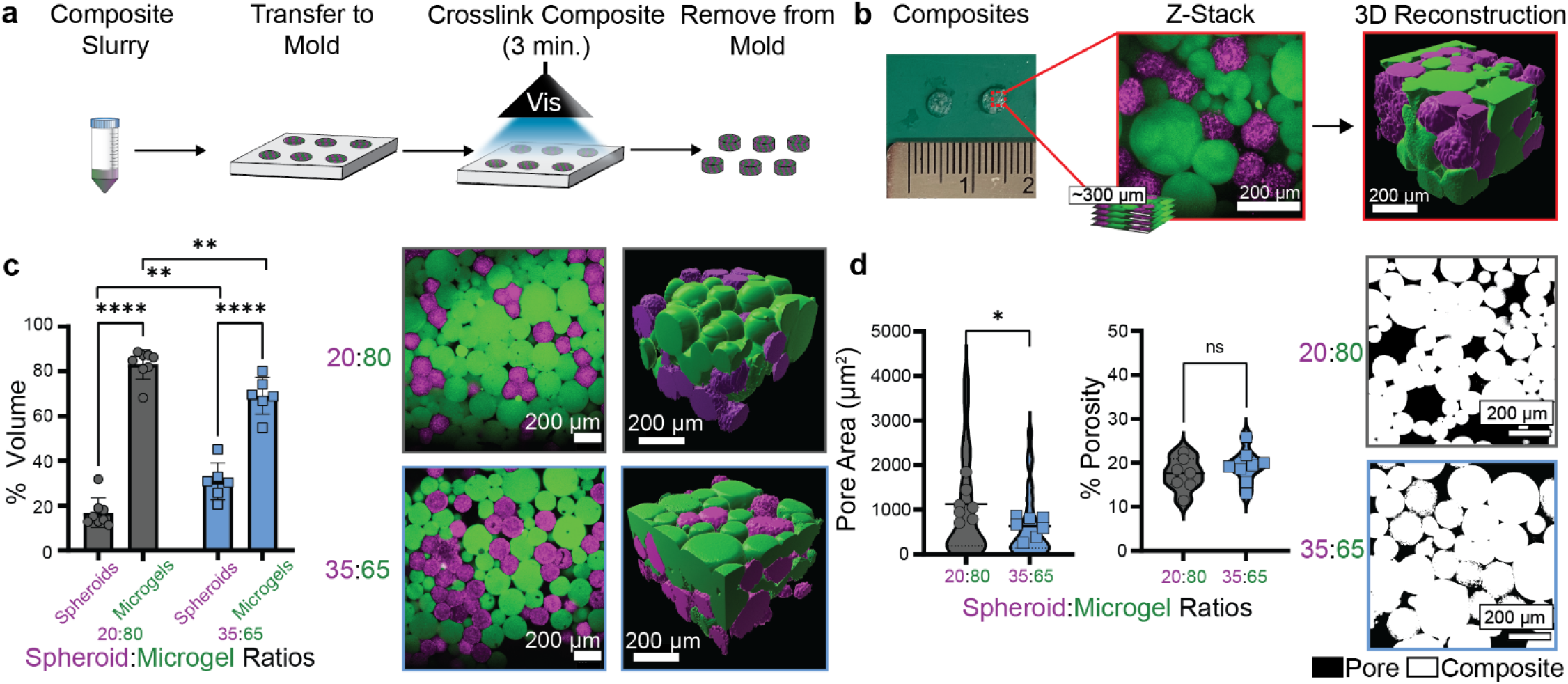
*In vitro* granular composite fabrication and characterization. **a**, Schematic of granular composite fabrication workflow. Composite slurries are transferred into 3D printed molds (Ø = 4 mm, H= 2 mm), exposed to visible light (20 mW/cm^2^) for 3 min. to crosslink microgels, and then removed from molds. **b**, Schematic of granular composite 3D reconstruction workflow for volume validation. Granular composites are formed with MSC spheroids stained with CellTracker Red and NorHA microgels incorporating FITC-dextran for visualization and processed through confocal microscopy (∼300 µm z-stack height and reconstruction via Imaris microscopy software) through RapiClear® tissue clearing; scale bars: 200 µm. **c**, Quantification of granular composite % total volume for microgel and spheroid components. n=8 (20:80), n=6 (35:65) composites from 2 biologically independent experiments, mean ± s.d. Representative images of 20:80 and 35:65 granular composites and representative 3D reconstructions; scale bars: 200µm. **d**, Quantification of pore area and % porosity for 20:80 and 35:65 granular composites. n=7 composites from 2 biologically independent experiments, mean ± s.d, ROUT method (Q = 1) determined outliers for pore area and % porosity measurements. Representative confocal slices of granular composites (white) with their porosity highlighted in (black); scale bars: 200 µm.

### Granular hydrogels support MSC chondrogenesis and cartilage formation

As 20:80 and 35:65 spheroid to microgel volume ratio granular composites are both injectable and stable immediately after interparticle crosslinking, these formulations are selected for culture in the presence of chondrogenic medium for up to 56 days (Fig. 4a). Given previous reports of spheroid cultures collapsing over time and losing shape fidelity, monitoring of granular composite diameter and shape is used to ensure stability of the constructs with culture (Fig. 4b). This indicates that the NorHA microgels provide enough mechanical support to prevent MSC spheroids from collapsing during fusion. Gene expression analysis via qPCR is next used to monitor MSC chondrogenesis, as well as evidence of fibrocartilage (*COL1A1*) and hypertrophic (*COL10A1*) markers (Supplemental Fig. 7a). Over 14 days of culture, granular composites increase expression of chondrogenic markers (*SOX9, ACAN, COL2A1*) hundreds to thousands in fold change when normalized to undifferentiated MSCs, while fibrocartilage and hypertrophic markers increase only minimally, particularly when compared to changes in chondrogenic markers. All genes apart from *COL1A1*, which is significantly increased in the 35:65 group, are not influenced by the tested spheroid to microgel volume ratios.

To evaluate the production of cartilage tissue, constructs are cultured for up to 56 days and the biochemical content, biomechanical properties, and histology are monitored. Over time, increases in dsDNA levels remain insignificant in both 20:80 and 35:65 volume ratios, indicating no drastic loss in cell viability; however, by D56, the 35:65 group has significantly higher dsDNA content compared to the 20:80 group, most likely due to the initial cell density difference (Fig. 5ci). Regarding ECM content, both groups generally exhibit significant increases in sGAG and collagen over time, but the 35:65 group exhibits ∼50-70% more ECM by D56 over the 20:80 group (Fig. 5cii, iii). We next determine the compressive properties of granular composites via uniaxial compression testing. At D1, the 20:80 granular composites exhibit ∼66% greater modulus than the 35:65 group, due to the increased microgel volume. Over time, both groups exhibit increases in compressive moduli until D56, where the 35:65 group (∼600 kPa) is significantly greater by ∼20% than the 20:80 group (∼500 kPa). These results are consistent with the observed increases in ECM content and suggest that increased mechanics relate to greater ECM deposition. The compressive moduli values are quite high for engineered cartilage tissue, particularly with the use of adult MSCs, likely due to enhanced early cell-cell contacts due to use of spheroids and the ability to elaborate a matrix due to the granular structure. Additionally, the values measured for cartilage formed with granular composites at D56 are within the range previously reported for native hyaline cartilage moduli, specifically 0.1-1.6 MPa.^33^

**Fig. 5.**
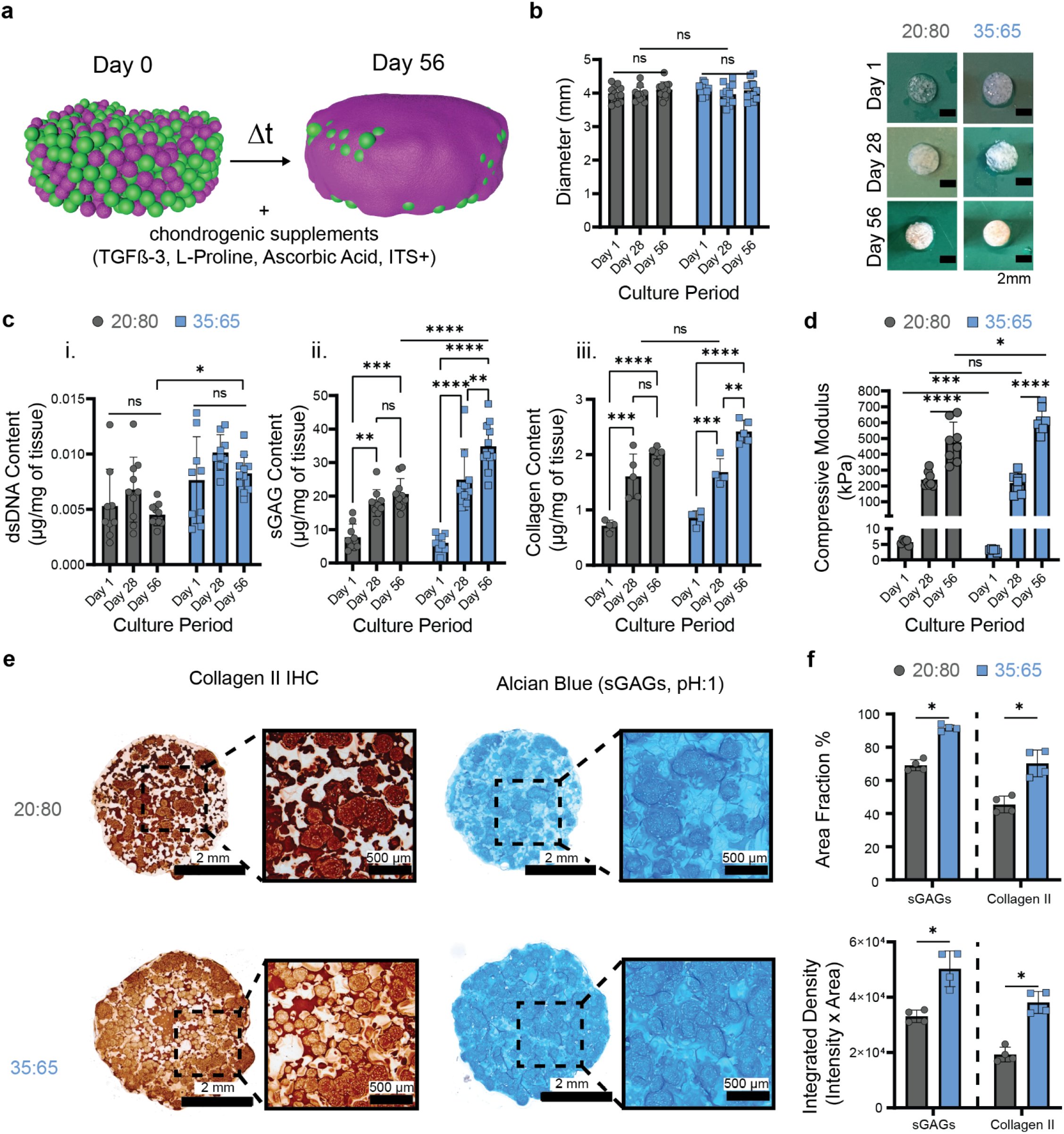
Long-term chondrogenic culture of granular composites. **a**, Schematic of granular composites, where spheroid fusion and growth results in cartilage tissue over time when cultured in the presence of chondrogenic factors. **b**, Quantification of diameters (left) and images (right) of granular composites with culture for 20:80 and 35:65 spheroid to microgel volume ratios. scale bars: 2 mm; n=12 composites from 2 biologically independent experiments; mean ± s.d. **c**, Quantification of (i) dsDNA, (ii) sGAG, and (iii) collagen contents of granular composites with culture for 20:80 and 35:65 spheroid to microgel volume ratios. dsDNA/sGAG: n= 9 (Day 1, Day 28: 35:65), n= 10 (Day 28, 56), n=11 (Day 56, 35:65) composites from 2 biologically independent experiments; mean ± s.d. Collagen: n=4 (Day 1: 20:80; Day 1, 28: 35:65), n=5 (Day 28, 56: 20:80; Day 56: 35:65) composites from 1 biologically independent experiment; mean ± s.d. **d**, Uniaxial compression values of granular composites with culture for 20:80 and 35:65 spheroid to microgel volume ratios. n= 5 (Day 1), n= 8 (Day 28, 56) from 2 biologically independent experiments; mean ± s.d. **e**, Representative histological images of granular composites for 20:80 and 35:65 spheroid to microgel volume ratios at day 56 stained for Alcian blue (pH:1, sGAG) and collagen II (IHC); scale bars: 2 mm; inset scale bars: 500 μm. **f**, Quantification of area fraction % and integrated density (intensity * area) of sGAG and collagen II for granular composites cultured at day 56. n= 4 independent composites; mean ± s.d.

To further assess ECM deposition and distribution within granular composites, sGAGs and collagens are stained at D28 and D56 (Fig 5e.; Supplemental Fig. 7b). Quantification of percent area fraction demonstrates that the 35:65 group deposits increased areas of sGAGs and collagen II by ∼35-50%, meaning less void spaces without matrix, and ∼35-100% more sGAGs and collagen II intensity, meaning greater matrix deposition throughout (Fig. 5f). These observations again are consistent with the measured levels of ECM and the greater microgel amounts introduced into the 20:80 constructs that can limit ECM distribution. To ensure a high balance of collagen II/I, often used to indicate a hyaline cartilage, granular composites are stained for collagen I and demonstrate very little staining at either D28 or D56 (Supplemental Fig. 7b).

### Chondrocytes enhance MSC-based cartilage tissue formation in composites

MSC therapeutic efficacy and potential for chondrogenesis diminishes with age;^34^ however, recent studies demonstrate that the addition of CHs to adult MSCs through co-cultures improves MSC chondrogenesis and ECM production.^23,24^ To investigate whether co-cultures within spheroids of our granular composite approach also improves cartilage formation, we introduce spheroids of a 4:1 MSC:CH ratio into the system based on previous reports on the chondrogenic potential of this ratio and our observation that this does not alter initial spheroid size (Fig. 6a, b).^23,24,35^ As the 35:65 group produces superior cartilage tissue, we fabricate our co-culture granular composites at this ratio and expose them to chondrogenic medium for 56 days. Co-culture granular composites significantly increase in size over this culture, when compared to the 35:65 MSC only granular composite control that does not (Fig. 6c). Additionally, the dsDNA and sGAG contents significantly increase over time and with the co-cultures, which could explain the size increase in cartilage constructs (Fig. 6di, ii). Interestingly, the collagen content increases over time in both groups, but is not statistically significant between the co-culture group and control (Fig. 6diii). Importantly, the compressive modulus is ∼30% greater in the co-culture composite (∼650 kPa) compared to the MSC only group at D56 when donor and passage matched MSCs are used (Fig. 6e).

**Fig. 6.**
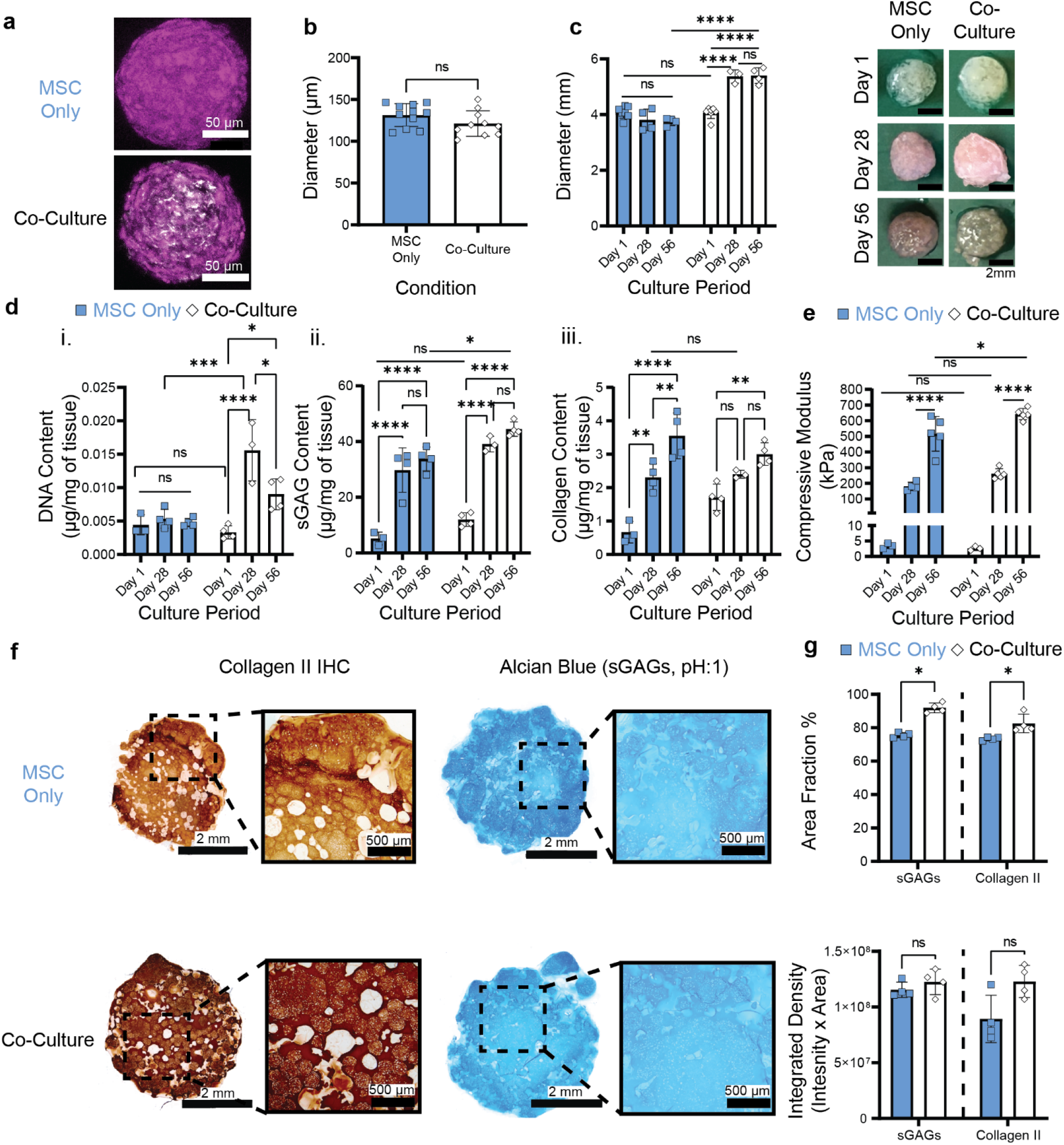
Long-term granular composite cultures are enhanced by chondrocytes. **a**, Representative images of MSC only and 4:1 co-culture spheroids; MSC (magenta), chondrocyte (white); scale bars: 50µm. **b**, Quantification of MSC only and co-culture spheroid diameter after 2 days condensation. n=11 (MSC only), n=10 (co-culture) spheroids from 1 biologically independent experiment; mean ± s.d. **c**, Quantification of granular composite diameter over varying culture periods and co-culture conditions. 35:65 MSC only (Blue), co-culture (White) composites. n=7 (Day 1), n=4 (Day 28,56; MSC only), n=4 (Day 56, co-culture), n=3 (Day 28, co-culture) from 1 biologically independent experiment; mean ± s.d. **d**, Quantification of biochemical content of granular composites at varying culture periods and co-culture conditions, including (i) dsDNA content; n=4 (Day 28, 56; MSC only), n=3 (Day 1; MSC only), n=4 (Day 1, 56; co-culture), n=3 (Day 28; co-culture) composites from 1 biologically independent experiment; mean ± s.d. (ii) sGAG content; n=4 (Day 28, 56; MSC only), n=3 (Day 1; MSC only), n=4 (Day 1, 56; co-culture), n=3 (Day 28; co-culture) composites from 1 biologically independent experiment; mean ± s.d. (iii) collagen content; n=4 (Day 28, 56; MSC only), n=3 (Day 1; MSC only), n=4 (Day 1, 56; co-culture), n=3 (Day 28; co-culture) composites from 1 biologically independent experiment; mean ± s.d. **e**, Uniaxial compression values of granular composites over varying culture periods and co-culture conditions. n=3 (Day 1), n=4 (Day 28; MSC only), n=5 (Day 56; MSC only) n=5 (Day 28, 56; co-culture) composites from 1 biologically independent experiment; mean ± s.d. **f**, Representative histological images of granular composites of varying co-culture conditions at D56 stained for Alcian blue (pH:1, sGAG) and collagen II (IHC); scale bars: 2 mm; inset scale bars: 500 μm. **g**, Quantification of area fraction % and integrated density (intensity * area) of sGAG and collagen II for granular composites at D56. n= 4 independent composites; mean ± s.d.

Cartilaginous ECM production, including sGAG, chondroitin sulfate, and collagen II, are abundant in both groups at D28 and D56 (Fig. 6f, Supplemental Fig. 8). Upon histological quantification, co-culture granular composites have ∼5-20% increase in percent area fraction for both sGAG and collagen II stains compared to MSC only control; however, similar integrated densities are observed across groups (Fig. 6g). Collagen I staining is again limited in both co-culture and MSC only groups at both D28 and D56 (Supplemental Fig. 8).

### Outlook

We demonstrate an innovative strategy where MSC spheroids and NorHA microgels are combined into granular composites for engineering cartilage tissue. This approach leverages desirable features of both spheroids and granular hydrogels, such as injectability through shear-thinning and self-healing properties of granular media, stability through interparticle crosslinking of microgels, cell-cell contacts to promote MSC chondrogenesis, and the formation of cartilage tissue with culture due to spheroid fusion and growth. Simulated mixing of granular composites reveals that the initial spheroid to microgel volume ratio influences the connectivity of components within granular composites, which is supported through experimental testing of compositions with high microgel interconnectivity that result in rapid initial construct stabilization and shape maintenance with culture. These granular composites possess ∼20% porosity initially, which fills with cartilage matrix during culture and results in mechanical properties that are greater than 200-fold from initial properties and are similar to those of native hyaline cartilage. Of note, microgels remain after culture and are only partially filled with ECM, providing future opportunities to enhance microgel degradation to further support ECM distribution, as others have observed in hydrogels.^36^ We also demonstrate that the addition of CHs to the spheroids is one approach to further improve formed cartilage properties; however, other approaches such as local growth factor delivery can be utilized to improve upon the modular design. One limitation of the study is that the granular composites are not implemented yet in cartilage defect models, which would likely require large-animal models to replicate the size-scales of clinical tissue defects.

When comparing to prior work in cartilage tissue engineering, there are numerous advantages to this approach. The shear-thinning and self-healing properties of the granular composites enables injectability, which is missing with many biomaterial and non-biomaterial approaches, such as with non-viscous hydrogel precursors or cell suspensions that may disperse prior to crosslinking. Although there are other examples of injectable materials that could be retained within defects, such as supramolecular assemblies, these systems lack a macroporous structure that may limit cell-cell contacts and ECM deposition.^37^ Additionally, prior work with MSC spheroids exhibit comparable mechanical properties to our approach, but the lack of injectability and limited initial construct stabilization will limit clinical translation.^31,38^ Although construct stability with spheroids can be introduced through seeding on macroporous lattice scaffolds, this approach is again not injectable and suffers from poor spheroid seeding and uniformity.^39–42^ MSCs may also be encapsulated directly within microgels, supporting injectability, but this often results in fibrocartilage production and limited cell-cell contacts.^43–45^ Together, it becomes apparent that this new granular composite approach for engineered cartilage tissue is very promising to advance the field.

## Methods

### Materials and Instruments

Sodium hyaluronate (HA, mol. wt. = 66 kDa) was purchased from Lifecore Biomedical (Chaska, MN), lithium phenyl-2,4,6trimethylbenzoylphosphinate (LAP) was purchased from Colorado Photopolymer Solutions (Boulder, CO), RapiClear® 1.49 (RC149001) was purchased from SunJin Lab Co. (Taiwan), and all immunohistochemistry supplies were purchased from Agilent (Santa Clara, CA) except for DAB chromogen stains (Sigma). Unless otherwise specified, all other reagents were purchased from Sigma-Aldrich and Fischer Scientific. Confocal microscopy: Leica SP5 II microscope. Rheology: TA Instruments AR2000ex. Uniaxial compression testing: TA Instruments Q800 DMA. Fluorescence intensity: Tecan Infinite 200 Pro with 480 nm excitation and 520 nm emission wavelengths for Picogreen assay, absorbance at 525 nm for sGAG DMMB assay, and absorbance of 560 nm for hydroxyproline assay. Histology imaging: Nikon SMZ18 stereoscope. PCR and qPCR: BIORAD CFX 96 real-time system.

### Polymer synthesis and characterization

Norbornene-modified HA (NorHA) was synthesized as previously described.^41^ Briefly, sodium HA was dissolved in DI water and mixed with Dowex 50W x 200 proton exchange resin (3:1 ratio, resin:HA) for 2 hrs., titrated to pH ∼7.02-7.05 with tetrabutylammonium hydroxide (0.2 M), frozen and lyophilized. HA-TBA methylation peak was determined, normalized to the methyl peak on HA (δ∼2.0-1.8) and measured (δ∼0.7-.9; 17.21) via ^1^H-NMR (Supplemental Fig. 10a). HA-TBA was modified with norbornene through benzotriazole-1-yl-oxy-tris-(dimethylamino)-phosphonium hexafluorophosphate (BOP) coupling. Briefly, HA-TBA and 5-norbornene-2-methylamine was dissolved with anhydrous dimethyl sulfoxide (DMSO), BOP was cannulated into the reagent solution and allowed to react for 2 hrs. under N_2_. The reaction was quenched with cold DI water, placed on dialysis (6-8 kDa mesh cutoff) for 2 days with DI water and salt and a subsequent 3 days with DI water, frozen and lyophilized. The extent of norbornene modification was determined via ^1^H-NMR to be ∼ 20-22% (δ∼5.8-6.2, 2H) of the disaccharide repeat units of HA when normalized to the methyl peak on HA (Supplemental Fig. 10b).

### Microgel fabrication and size characterization

NorHA microgels were fabricated via water-in-oil emulsion. A 3% NorHA precursor solution was prepared with 0.05% I2959, 10 mM dithiothreitol (DTT), and 1x phosphate buffer saline (PBS), added to oil phase (98% light mineral oil/ 2% Span 80) dropwise, allowed to stir for 30 sec., and crosslinked with a UV (320-390 nm) omnicure lamp at 20 mW cm^-2^ for 30 min. Oil phase stir speed was varied to obtain varying size distributions. Microgels were washed by discarding oil phase, soaking in 1% tween 20 solution for 10 min., and washing with 1x PBS, which was repeated 5x. Microgels were collected, centrifuged at 15,000 x g for 5 min. and stored at 4 °C before use in experiments. Fluorescein isothiocyanate (FITC)-dextran (MW: 2 million Da) was added into the NorHA precursor solution to allow visualization of microgels for size characterization. Microgels were visualized with a fluorescent microscope, thresholded, and sizes (ferret diameter) were measured via analyze particle function (FIJI).

### Microwell fabrication

Microwell pyramid arrays were fabricated from a poly(dimethyl)siloxane (PDMS) molding process to generate spheroids. PDMS was mixed in a 10:1 polymer to crosslinker ratio, degassed, added into commercial microwell plates (StemCell Technologies, AggreWell™ 400) and placed into an oven at 80°C for 2 hrs. to form a negative mold. Negative molds were removed, surface treated with Trichloro(1H,1H,2H,2H-perfluorooctyl)silane for 45 min. via vapor deposition after O_2_ plasma activation with plasma etcher (Plasma Etch) for 3 min. Positive PDMS molds were created by adding uncured PDMS into each 6 well, degassing, positioning the negative PDMS mold, and curing in the oven at 80 °C for 2 hrs. PDMS positive molds were soaked in isopropyl alcohol for 1 hr., left in DI water overnight, and sterilized under a UV germicidal tissue culture hood for 25 min. Each well was rinsed with 1x PBS and an anti-adherence rinsing solution (StemCell Technologies, Aggrewell™) was added into each well. Plates were centrifuged at 2000 x g for 5 min to remove air bubbles. The wells were aspirated, rinsed with basal medium, and placed in an incubator (37 °C/ 5% CO_2_) with chondrogenic medium until cell seeding.

### Cell isolation and expansion

Cells were obtained from adult Yucatan mini-pigs (porcine). Femoral bone marrow was extracted and MSCs were isolated via plastic adherence during culture in Dulbecco’s modified eagle medium (DMEM) with 10% fetal bovine serum (FBS), and 1% penicillin/streptomycin (P/S) according to previous literature.^46^ All donors were seeded at 5000 cells cm^-2^, with alpha modified eagle medium (α-MEM) supplemented with 10% FBS, 1% P/S, and fibroblast growth factor (FGF, 1 ng mL^-1^). Articular chondrocytes (CH) were isolated from cartilage on the femoral condyle head. Cartilage pieces were obtained with a scalpel, minced, and placed into adherent tissue culture dishes with DMEM, collagenase II (∼382 U mL^-1^), 2% P/S, and incubated for 2 days. A 50 μm cell strainer was used to collect the isolated CHs and remove unwanted tissue debris. CHs were plated into adherent tissue culture plates at 70,000 cells cm^-2^ up to passage 2 to preserve chondrogenic phenotype in DMEM, 10% FBS and 1% P/S.

### Spheroid formation, size and viability characterization, and culture

#### Spheroid Formation

Porcine MSCs were rinsed with 1x PBS, trypsinized, and resuspended in chondrogenic medium at varying densities, such that the total volume in each well was 5 mL. Cells were allowed 48 hrs. to condensate into spheroids. Chondrogenic medium consisted of DMEM + Glutamax, 10% FBS, 1% P/S, 1% ITS, 1 × 10^−3^ M sodium pyruvate, 50 µg ml^-1^ ascorbic acid 2-phosphate, 40 µg ml^-1^ L-proline, 1 × 10^−7^ M dexamethasone, and 10 ng ml^-1^ transforming growth factor-β3 (TGF-β3). Co-culture spheroids were formed identically except the cell suspension was comprised of MSCs and CHs in a 4 to 1 ratio. Spheroid size was determined after condensation via brightfield microscopy. Co-culture spheroids were visualized via confocal microscopy by staining CHs with CellTracker Red CMTPX (Invitrogen, 12.5 µM) and MSCs with CellTracker Green CMFDA (Invitrogen, 15 µM) for 45 min. at 37°C prior to spheroid fabrication. *Spheroid Viability*. Spheroid viability was evaluated at days 1 and 7 via Calcein AM (Invitrogen, 2 µM) and ethidium homodimer (Invitrogen, 4 µM) by staining for 3 hrs. at 4 °C to allow diffusion throughout the spheroid. Cell viability was quantified from confocal stacks and reported as % live cell area, which was calculated by the ratio of Calcein AM stained area to the combined Calcein AM and ethidium homodimer stained area. *Spheroid Culture*. MSCs were seeded at 1000 cells spheroid^-1^ in chondrogenic medium and spheroid growth were monitored (ferrets diameter) for 28 days of culture. Chondrogenic medium was replenished every 2-3 days, imaged with a brightfield phase contract microscope every 7 days, and quantified via analyze particle function (FIJI). Spheroid volume was calculated assuming a perfect sphere.

### Immunofluorescent staining

Spheroids were stained at varying culture periods (day 1, 7, 28) by fixing in 10% formalin at 4 °C overnight, blocking with 1% bovine serum albumin (BSA), and washing with 1x PBS. Samples were blocked, and anti-chondroitin sulfate mouse monoclonal antibodies (Sigma SAB4200696, 1:200) and anti-collagen II mouse monoclonal antibodies (DSHB II-II6B3, 1:100) were introduced overnight at 4 °C. Secondary antibody staining was performed using goat anti-mouse polyclonal IgG (H+L) AlexaFluor™ 647 (Invitrogen A-20990, 1:1000) for 2 hrs. at 25 °C. Samples were washed, permeabilized in 2% triton X-100 overnight at 25 °C, and cleared in 2 hrs. of adding RapiClear® solution. Confocal z-stacks (∼150 µm thickness) were taken to capture chondroitin sulfate and collagen II matrix deposition.

### Granular composite fabrication and long-term chondrogenic cultures

MSC only or co-culture spheroids were disrupted from the microwells via mixing with pipette, washed with 1x PBS, and transferred to a conical tube on ice. Spheroid suspensions were centrifuged at 300 x g for 20 sec. and NorHA microgels were centrifuged at 15,000 x g for 5 min. to determine total volume. 2.5 mM DTT and 0.05% LAP photoinitiator was added to the microgels to allow for interparticle crosslinking, vortexed, and centrifuged at 15,000 x g for 5 min. The spheroids and microgels were then added together at varying volume ratios, manually mixed with a spatula, and transferred into 3D printed molds (Ø: 4 mm, H: 2 mm) to photocrosslink with a visible light (400-500 nm) omnicure lamp at 20 mW cm^-2^ for 3 min. Granular composites were removed from 3D printed molds and placed within a 24-well non-adherent TC plate with chondrogenic medium. Long-term chondrogenic cultures were maintained up to 56 days and the medium was replenished every 2-3 days. Gross granular composite diameters were manually measured at varying culture periods (day 1, 28, 56) via measure tool (FIJI).

### Granular composite 3D reconstructions, volume, and porosity analysis

To image granular composites, MSCs were stained with CellTracker Red CMTPX and (FITC)-dextran was added into NorHA microgels as previously described above. Granular composites were fabricated, immediately fixed with 10% formalin overnight at 4 °C, permeabilized with 2% triton X-100 overnight at 25 °C and cleared with RapiClear® solution for 2 hrs. before imaging. Confocal z-stacks (∼300 µm) were imaged, and 3D reconstructions were created in Imaris 9.8 Microscopy Software. Total volume of each phase was exported and used to validate % total volume of the granular composite components. % Total volume is reported as volume of microgels or spheroids normalized to the total volume of microgels and spheroids * 100. Pore volume was excluded from this calculation. Porosity area and % porosity of granular composites was manually measured via 2D slices from confocal z-stacks (FIJI). % Porosity is reported as the % area of binary pores within the slice normalized to the total % area of both granular composite and pores.

### Mechanical testing

#### Rheological Testing

Granular composites were transferred onto the base platform while a 20 mm acrylic parallel plate was lowered to a gap height of 1 mm. Oscillatory rheological time sweeps (1% strain, 1 Hz) were performed to obtain the average storage (G’) and loss (G”) moduli. Oscillatory rheological strain sweeps (1-100% strain, 1 Hz) were performed to demonstrate shear-thinning properties. Oscillatory strain on/off sweep (500%/0% strain, 1 Hz) were performed to demonstrate self-healing properties. Granular microgels or spheroids were tested with oscillatory rheological time sweeps. *Uniaxial Compression Testing*. Granular hydrogels and granular composites of varying volume ratios were crosslinked into cylindrical disks (Ø: 4 mm, H: 2 mm). Granular hydrogels crosslinked at varying exposure times (0.5, 1, 3, 5 min.; 400-500 nm 20 mW cm^-2^) were tested to achieve an optimal crosslink time. Granular composites were tested at varying culture periods (1, 28, 56 days). Granular hydrogels and day 1 granular composites were compressed (0.05 N min^-1^) with a preload of 0.001 N. Day 28 and 56 granular composites were compressed to (0.5 N min^-1^) with a preload 0.05 N. Compressive moduli were calculated and reported as the stress-strain slope between 10-20% strain.

### Biochemical Content Analysis

#### Composite Digestion

Granular composites were digested as previously described with slight modifications.^41^ Briefly, samples were homogenized in 100 µg mL^-1^ of proteinase K, and 1 mg mL^-1^ hyaluronidase (750-3000 U mL^-1^) RNAse free solution. Samples were homogenized (Fisher Scientific, Bead Mill 24; 3.55 m s^-1^; three 5 min. cycles) with a 5 mm S.S. bead and placed in a 65 °C water bath to digest for 24-48 hrs. *Picogreen dsDNA Biochemical Analysis*. Total dsDNA content was measured with a Picogreen dsDNA assay kit (Invitrogen, P7589) according to manufacturer’s protocol and previous literature.^47^ Samples were diluted 1:10 (100 µL) compared to standards to fit within the working curve. *Dimethylmethylene Blue (DMMB) sGAG Biochemical Analysis*. Total sGAG content was measured via DMMB colorimetric assay as previously published.^47^ Briefly, chondroitin sulfate (Fischer Scientific, AAJ6034106) standards were created and samples were diluted 1:20 compared to standards to fit within the working curve. *Hydroxyproline Collagen Biochemical Analysis*. Total collagen content was measured with a hydroxyproline assay kit (Abcam, ab222941) according to manufacturer’s protocol. Briefly, 10 N NaOH was added to each sample, heated to 120 °C for 1.5 hrs., and neutralized with 10 N HCl once samples reach 25 °C. Collagen content was calculated by hydroxyproline: collagen ratio 7.14:1 as previously reported in literature.^47^ All biochemical content is also reported as µg/construct in Supplemental Fig 9.

### Histology and Immunohistochemistry

#### Histology

Samples were fixed overnight at 4 °C in 10% formalin, dehydrated with ethanol (50-100 v/v%), embedded with high-melt paraffin wax, and sectioned (5 µm) prior to staining. All samples were deparaffinized with citrisolv detergent and dehydrated before staining. sGAG matrix deposition was visualized with Alcian blue staining (Newcomer Supply, 1% pH:1). Samples were mounted in a toluene-based mounting solution and sealed with clear nail polish. *Immunohistochemistry*. Samples were stained as previously described.^41^ Collagen I, II, and chondroitin sulfate were visualized via staining for anti-collagen type I mouse monoclonal antibodies (Sigma MAB3391, 1:100), anti-collagen II mouse monoclonal antibodies (DSHB II-II6B3, 1:100), anti-chondroitin sulfate mouse monoclonal antibodies (Sigma SAB4200696, 1:200) overnight at 4 °C., followed by biotinylated secondary antibody staining and Streptavidin HRP for 10 min. DAB chromogen staining (1:25, A to B chromogens) was used to visualize stains by incubating for 10 min at 25 °C (Millipore). Samples were mounted in an aqueous-based mounting solution and sealed with clear nail polish. Histology images were white-balance corrected and % area fraction and integrated density was calculated (FIJI).

### Gene expression analysis

#### RNA Isolation, Polymerase Chain Reaction (PCR), and quantitative PCR (qPCR)

Samples were flash frozen with liquid N_2_ and sample RNA was extracted on ice via TRIzol reagent method described previously with slight modifications.^48^ Briefly, samples were homogenized for 30 sec. at 3.55 m s^-1^ cycle speed in TRIzol reagent, followed by phase separation with chloroform and precipitation with isopropanol. RNA was washed 2x with ethanol and resuspended in 60 µL of RNAse-free water. Total RNA was reverse transcribed with a high-capacity cDNA reverse transcription kit (Applied Biosystems, 4368814) according to manufacturer’s instructions. qPCR was carried out using SYBR green master mix (Applied Biosystems, A25742) according to manufacturer’s protocols. Primer forward and reverse sequences are reported in Supplemental Table 1. qPCR was measured in duplicate and fold change was reported as ΔΔCT method normalizing to GAPDH housekeeping gene and then undifferentiated MSCs.

### Cinema4D connectivity simulations and analysis

#### Connectivity simulations

To simulate mixing of composites, microgel and spheroids were simulated as rigid body spheres (Ø: 0.14 cm) at a 1:1000 size scale to reduce computational processing power. The ratio of spheroids and microgels were varied while keeping the total particle number to 372 particles. Each particle was assigned a dynamic body tag to introduce rigid body dynamic physics for mixing simulations. Parameters were Bounce = 0%, Friction = 0%, Collision Noise = 30%. A cylindrical tube (IR: 1.75 cm, OR: 5 cm) and 2 planes (400 cm x 400 cm) were used to contain mixing simulation and were given collision body tags to introduce parameters: Bounce = 0%, Friction = 0% and Collision Noise = 30%. Simulations were run for 70 frames per second, baked, and the current state was converted to an object that was exported as a .stl file. To introduce microgel size distributions, a step mograph effector was created and a spline was matched to an experimental size distribution as demonstrated in Supplemental Figure 4. Volume ratios were maintained using an area and volume function plugin. To introduce spheroid aggregation, a force mograph effector was introduced to the spheroids. Parameters were Strength = 3 cm, Damping = 0.1, Falloff = Step, IR = 0.14 cm, and OR = 0.141cm. For all simulations, gravity dynamics were set to parameters: Time Scale = 100% and Gravity = 10 cm. *Simulation analysis*. Files were uploaded onto FIJI’s 3D Viewer and voxelized (512 × 512 pixels x 50 z-slices) into binary z-stacks that were altered with smoothing and fill holes binary functions (FIJI). Connectivity was analyzed with 3D Objects Counter (FIJI) that determines the number of connected objects within a 3D space. Connectivity is defined as average number of spheroids or microgels in “clusters” divided by the total number of spheroids or microgels within the composite (∼0 = no connectivity, 1 = fully interconnected). Simulations were repeated 5 times with different seeds for each composite to introduce variability.

### Statistical analysis

All statistical tests were performed in GraphPad Prism 9. Comparison between two groups were analyzed with a two-tailed Student’s T test (if normality is assumed) or a Mann-Whitney test (if normality is not assumed). Normality calculated with D’Agostino & Pearson test; α = 0.05. Comparison between groups > 2 were analyzed with one-way or two-way ANOVA and multiple comparisons between groups were analyzed with Tukey multiple comparisons test with an α = 0.05 and 0.95 confidence interval. Bar graph descriptors (mean ± standard deviation, s.d.) are displayed within the figure panel caption. Experimental n values and biological replicates are stated in figure captions. % Porosity and pore area outliers were removed with ROUT test (Q = 1) in Fig. 4d. All statistical test p, q, t, and DF values are reported in Supplemental Table 2.

## Acknowledgements

This work was supported by the National Institutes of Health (R01AR077362) and the National Science Foundation through the Center for Engineering Mechanobiology STC (CMMI: 15-48571). We would also like to thank Drs. Katrina Wisdom and Jonathan Galarraga for their contributions with pilot experiments and insightful conversations.

