## Supplemental Information for "Injectable MSC Spheroid and Microgel Granular Composites for Engineering Cartilage Tissue"

**Supplemental Figures:**

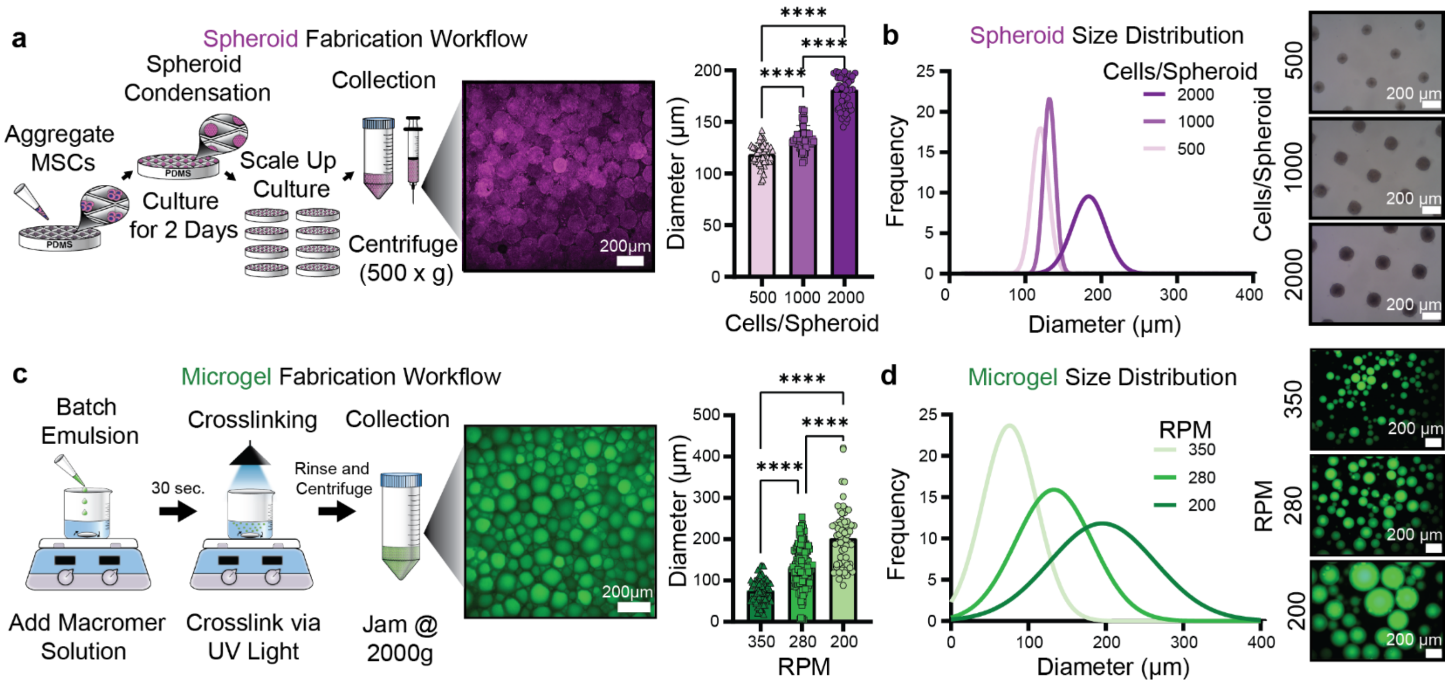


**Supplemental Figure 1 | Microgel and spheroid fabrication workflow, mean diameter, and size distribution analysis. a**, MSCs are seeded into PDMS microwell molds at varying cell seeding densities and allowed to condense for 2 days before collection. Molds are rinsed twice to remove all spheroids and are centrifuged to collect and measure diameter; scale bars: 200 μm. Quantification of mean spheroid diameter at varying cell seeding densities. n=50/condition from 3 biologically independent donors, mean ± s.d. RPM: revolutions per minute. **b**, Quantification of MSC spheroid size distribution (after the 2-day condensation period) at varying cells/spheroid seeding density with representative images of spheroids. n=50 spheroids/condition from 3 biologically independent donors; scale bars: 200 μm. **c**, NorHA microgels are fabricated via water-in-oil batch emulsion at varying emulsion spin speeds. NorHA prepolymer solution is added and allowed to mix for 30 sec. before crosslinking via UV light (20 mW/cm$^2$) for 30 min. Microgels are washed to remove residual oil and jammed to collect and measure diameters; scale bars: 200μm. Quantification of mean microgel diameters at varying batch emulsion speeds. n= 65 (200RPM) and 68 (350RPM) from 2 independent batch emulsions, mean ± s.d. **d**, Quantification of microgel size distribution fabricated via batch emulsion spun at varying RPM with representative images of NorHA microgels. n= 65 (200RPM), and 68 (350RPM) from 2 independent batch emulsions; scale bars: 200 μm. RPM: revolutions per minute.

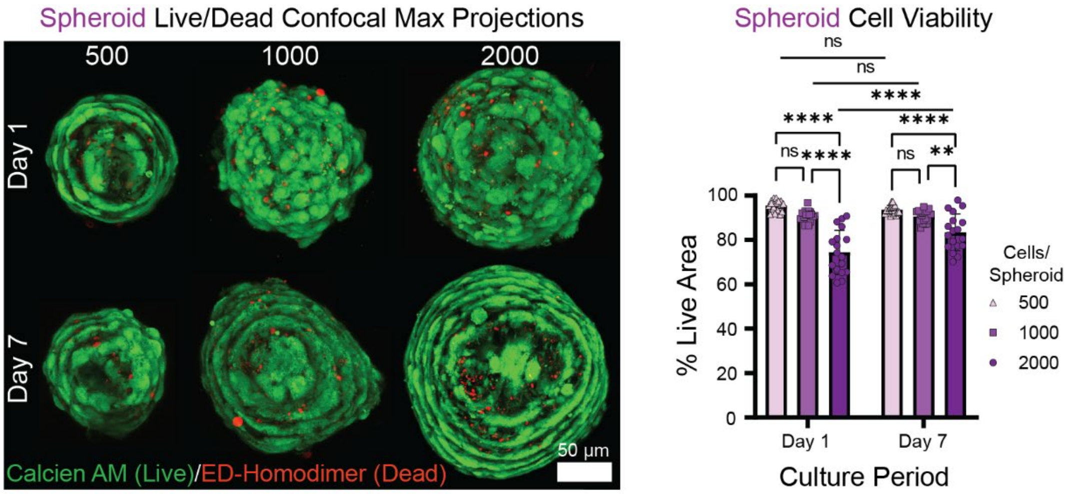


**Supplemental Figure 2 | MSC spheroid cell viability decreases as cells/spheroid and diameter increase.** Representative live (green)/dead(red) cell viability images of MSC spheroids of varying cell densities cultured for 1 or 7 days (after a 2-day condensation period); scale bars: 50 µm (**Left**). Quantification of % live area for varying MSC spheroid cell densities at 1 and 7 days of chondrogenic culture (**Right**). n=18 spheroids per cells/spheroid groups from 3 biologically independent donors, mean ± s.d.

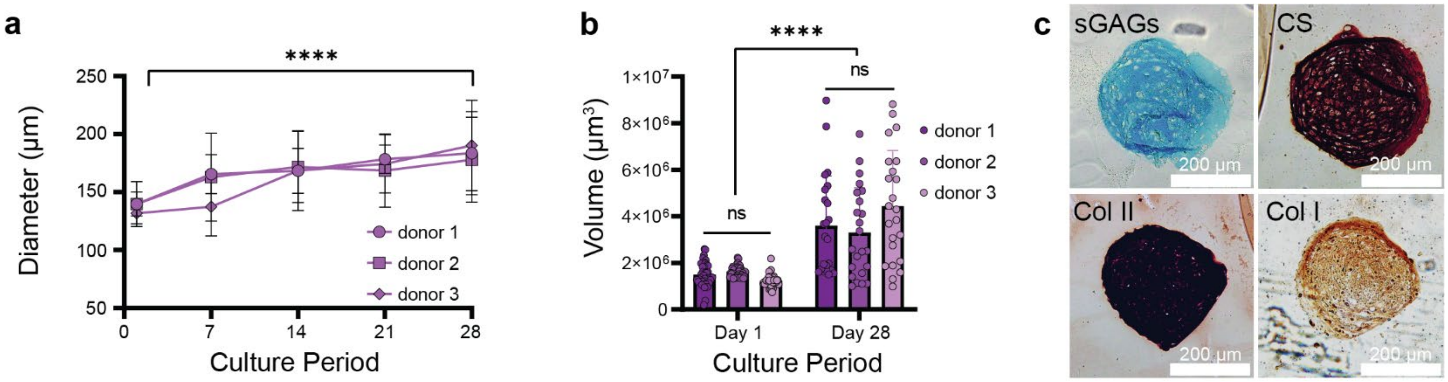


**Supplemental Figure 3 | Chondrogenic culture of individual MSC spheroids over time. a**, Quantification of MSC spheroid diameter growth over time. n= 37 spheroids (day 1), 51 (day 7), 45 (day 14), 19 (day 21), 23 (day 28) from 3 biologically independent donors, mean ± s.d. **b**, Volume measurements calculated via spheroid diameter values with the assumption of a perfect sphere ($V= 4/3\pi*r^3$). Spheroids grow 2-fold in volume compared to ~1.35-fold increase in spheroid diameter. n=37 (day 1), 23 (day 28) from 3 biologically independent donors, mean ± s.d. **c**, Representative day 28 MSC spheroid cultures in chondrogenic media with sGAGs (Alcian blue), chondroitin sulfate, collagen II and I (IHC) stains; scale bars: 200 µm.

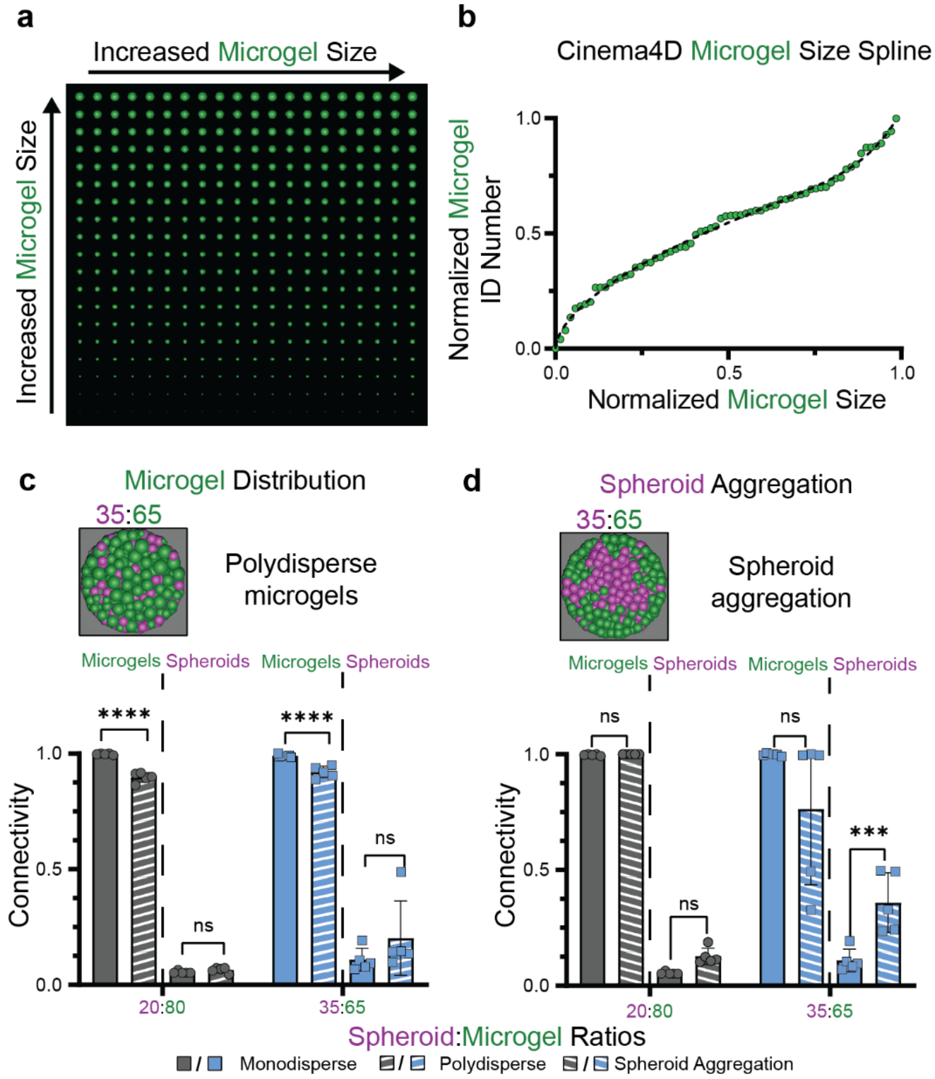


**Supplemental Figure 4 | Cinema4D microgel size distribution and spheroid aggregation simulations. a**, Representative image of a Cinema4D microgel size distribution array used to model granular composite connectivity for polydisperse conditions. **b**, Cinema4D spline curve imported into the software to introduce a microgel size distribution for polydisperse conditions that mimics experimental microgel size distribution (280RPM condition). n=68 microgels used to model spline. **c**, Quantification of granular composite connectivity that incorporates microgel size distribution into the simulation for 20:80 and 35:65 volume ratios. n=5 independent simulations, mean ± s.d. **d**, Quantification of granular composite connectivity that incorporates spheroid aggregation into the simulation for 20:80 and 35:65 volume ratios. n=5 independent simulations, mean ± s.d.

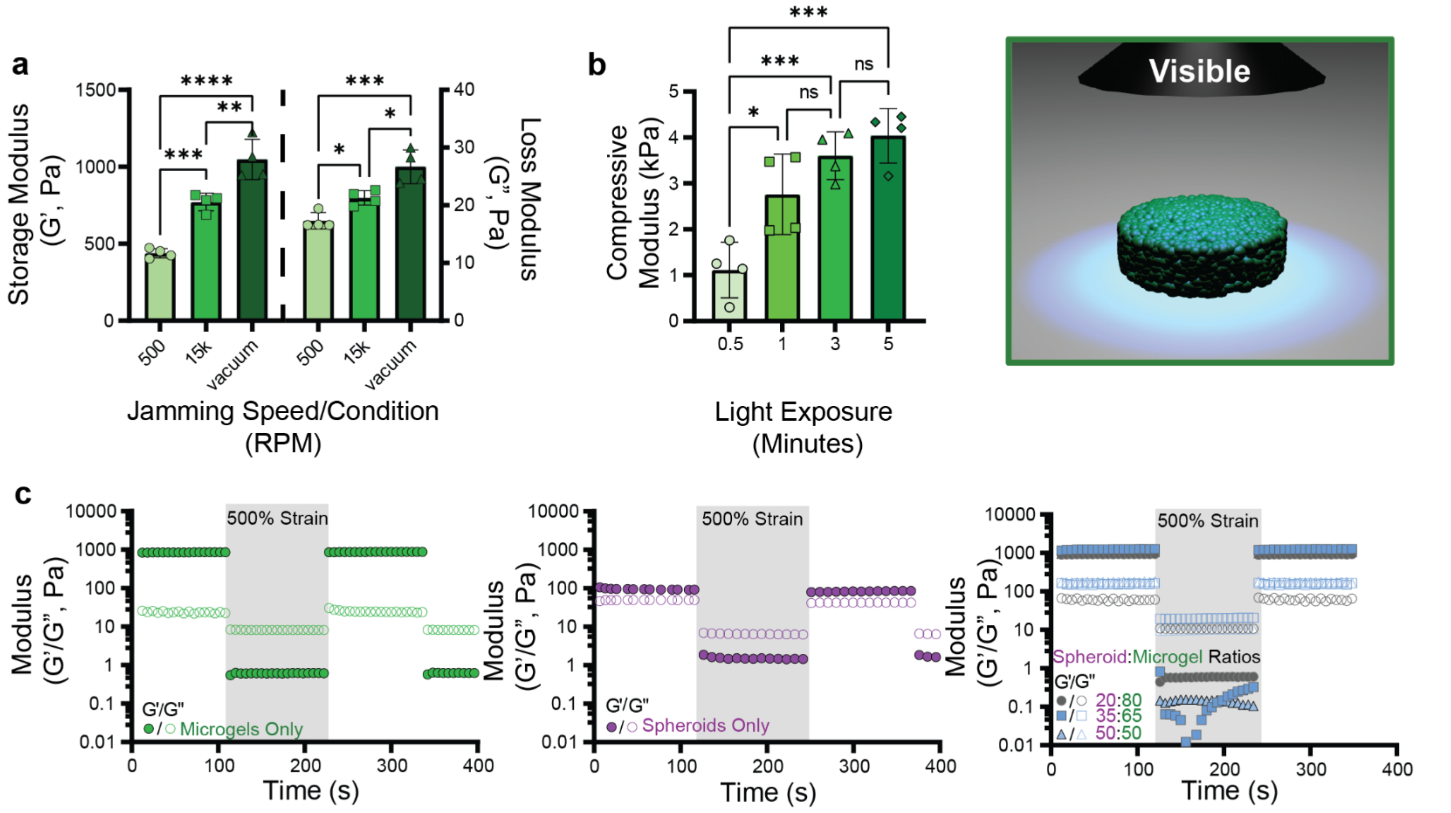


**Supplemental Figure 5 | Granular composite rheological characterization and post-crosslinking kinetics.** **a**, Quantification of granular hydrogels jammed at varying speeds and conditions. n=4 from independent rheological runs; mean ± s.d. **b**, Quantification of post-crosslinked granular hydrogels compressive modulus at varying visible light exposure times. n=4 independent hydrogel constructs; mean ± s.d. Schematic of granular hydrogel construct being post-crosslinked with visible light. **c**, Representative strain on/off (500%/1%) rheological plots for granular composite components and varying spheroid:microgel ratios. G' (storage modulus) is filled points; G" (loss modulus) is empty points. Shaded area: 500% strain, white area: 1% strain.

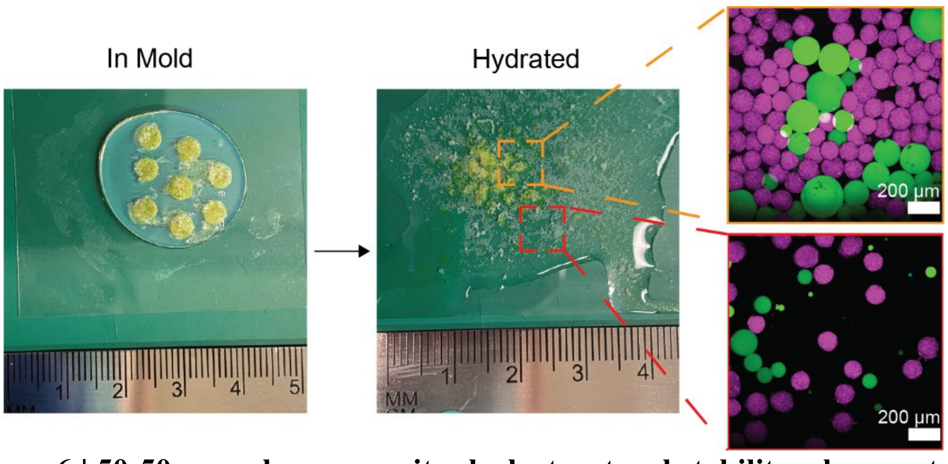


**Supplemental Figure 6 | 50:50 granular composites lack structural stability when post-crosslinked due to microgel connectivity.** Granular composites crosslinked with blue light (20 mW/cm$^2$) within molds, removed and hydrated with 1x PBS. 50:50 granular composites lack structural stability to stay together when hydrated, as visualized with macroscopic images and confocal imaging; scale bars; 200 µm.

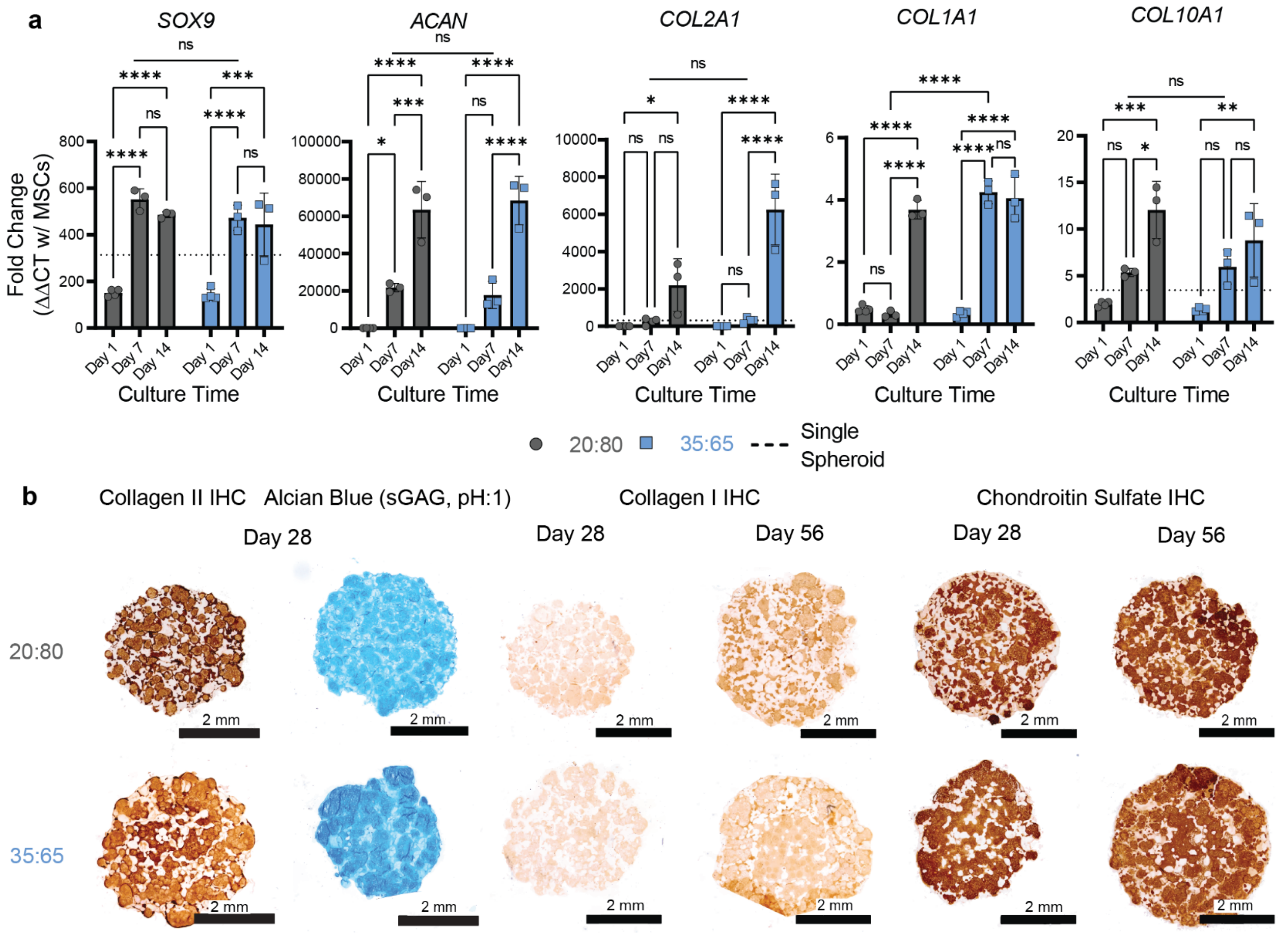


**Supplemental Figure 7 | Early timepoint qPCR for chondrogenesis of granular composites and IHC staining of granular composite slices at varying culture periods and volume ratios. a**, Quantification of qPCR chondrogenic genes of interest (SOX9, ACAN, COL2A1, COL1A1, TGFB3, COL10A1) via ΔΔCT method over varying culture periods and volume ratios. Fold change normalized to undifferentiated MSCs. n=4 (Day 1), n=3 (Day 7, 14) from independent granular composites. **b**, Representative histological slices of granular composites of varying volume ratios at Day 28 and 56 stained for collagen II (IHC), Alcian blue (pH:1, sGAG), collagen I (IHC) and chondroitin sulfate (IHC); scale bars: 2 mm.

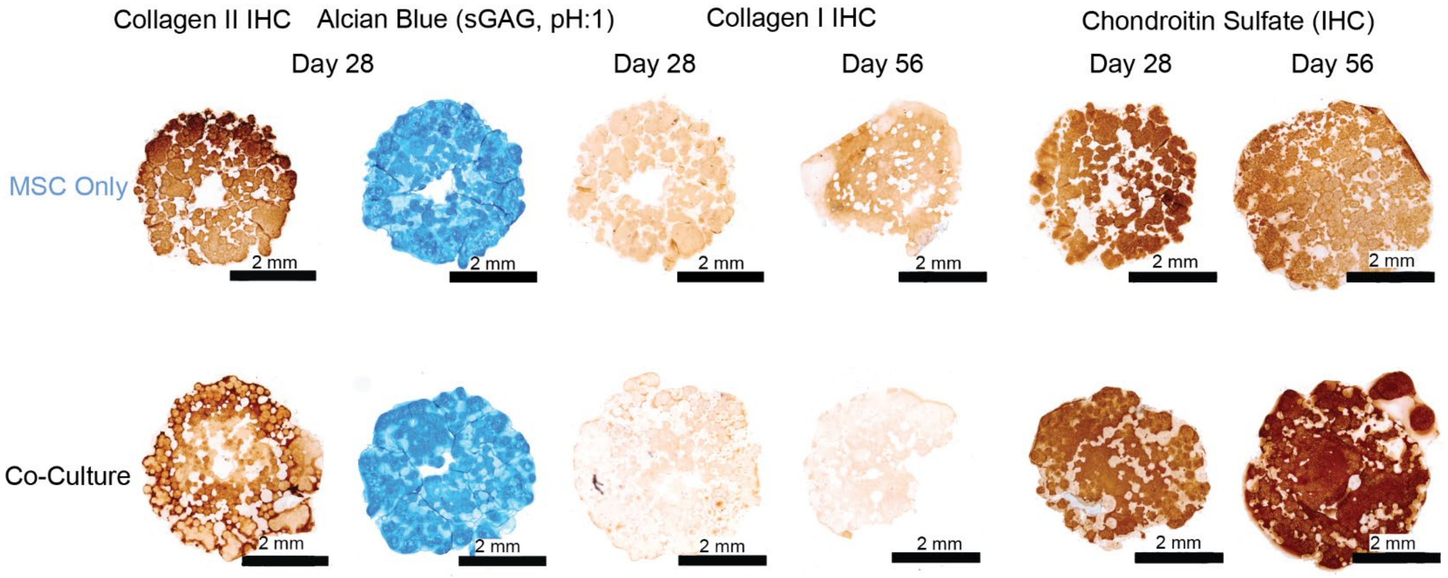


**Supplemental Figure 8 | IHC and histological staining of granular composites with varying culture periods and co-cultures conditions.** Granular composite histological slices of varying culture periods and co-culture conditions (MSC Only vs 4:1 co-culture) stained for collagen II, Alcian blue (sGAG, pH:1), collagen I and chondroitin sulfate; scale bars: 2 mm.

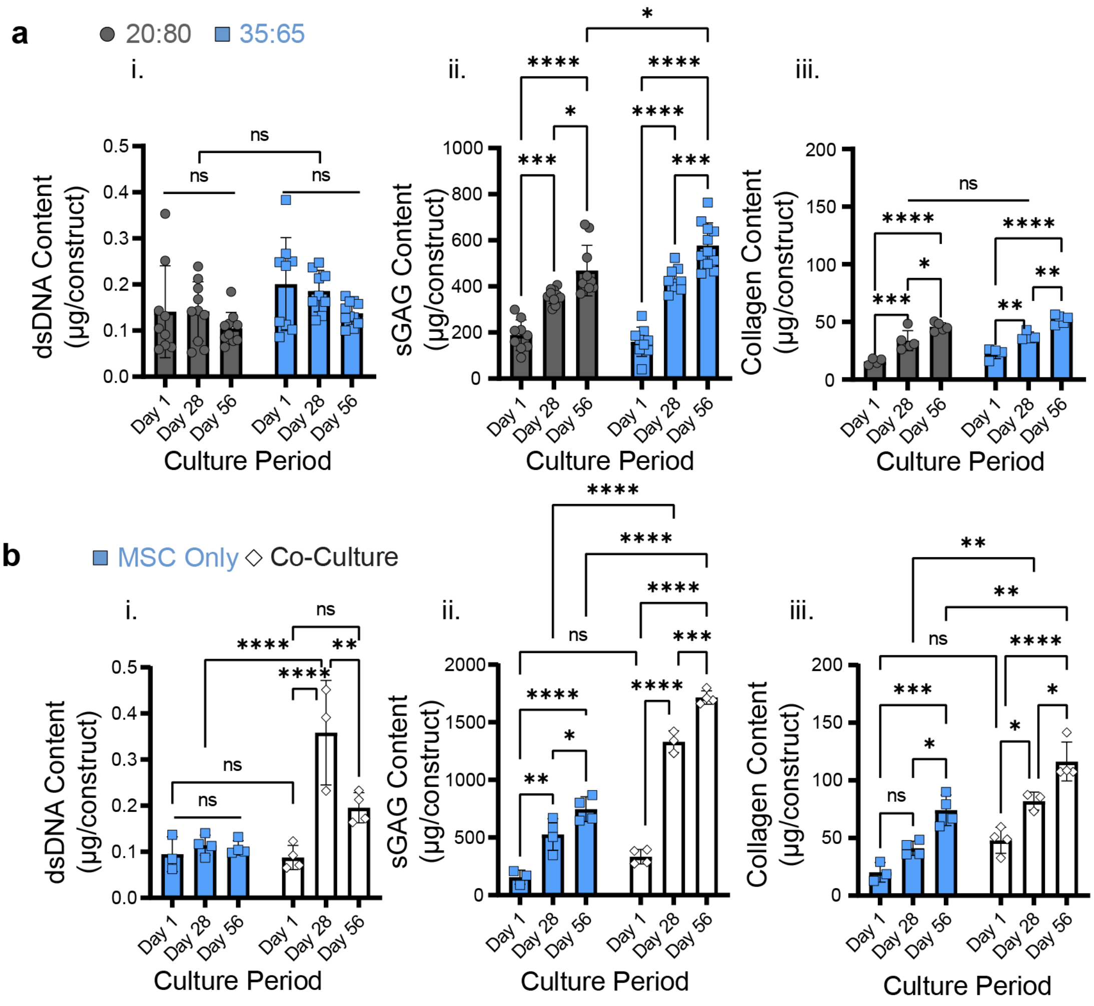


**Supplemental Figure 9 | Biochemical content of granular composites of varying volume ratios and co-culture conditions reported as µg per construct. a**, Quantification of biochemical content (DNA, sGAG, and Collagen) of granular composites at culture periods and varying volume ratios. (i) Quantification of dsDNA content of granular composites over varying culture periods and volume ratios. n= 9 (Day 1, Day 28: 35:65), n= 10 (Day 28, 56), n=11 (Day 56, 35:65) composites from 2 biologically independent experiments; mean ± s.d. (ii) Quantification of sGAG content of granular composites over varying culture periods and volume ratios. n= 9 (Day 1, 28: 35:65), n= 10 (Day 28, 56), n=11 (Day 56, 35:65) composites from 2 biologically independent experiments; mean ± s.d. (iii) Quantification of collagen content of granular composites over varying culture periods and volume ratios. n=4 (Day 1: 20:80; Day 1, 28: 35:65), n=5 (Day 28, 56: 20:80; Day 56: 35:65) composites; mean ± s.d. **b**, Quantification of biochemical content (DNA, sGAG, and Collagen) of granular composites at culture periods and varying co-culture conditions. (i) Quantification of dsDNA content of granular composites over varying culture periods and co-culture conditions. n=4 (Day 28, 56; MSC only), n=3 (Day 1; MSC only), n=4 (Day 1, 56; co-culture), n=3 (Day 28; co-culture) composites from 1 biologically independent experiment; mean ± s.d. (ii) Quantification of sGAG content of granular composites over varying culture periods and co-culture conditions. n=4 (Day 28, 56; MSC only), n=3 (Day 1; MSC only), n=4 (Day 1, 56; co-culture), n=3 (Day 28; co-culture) composites from 1 biologically independent experiment; mean ± s.d. (iii) Quantification of collagen content of granular composites over varying culture periods and co-culture conditions. n=4 (Day 28,

56; MSC only), n=3 (Day 1; MSC only), n=4 (Day 1, 56; co-culture), n=3 (Day 28; co-culture) composites from 1 biologically independent experiment; mean ± s.d.

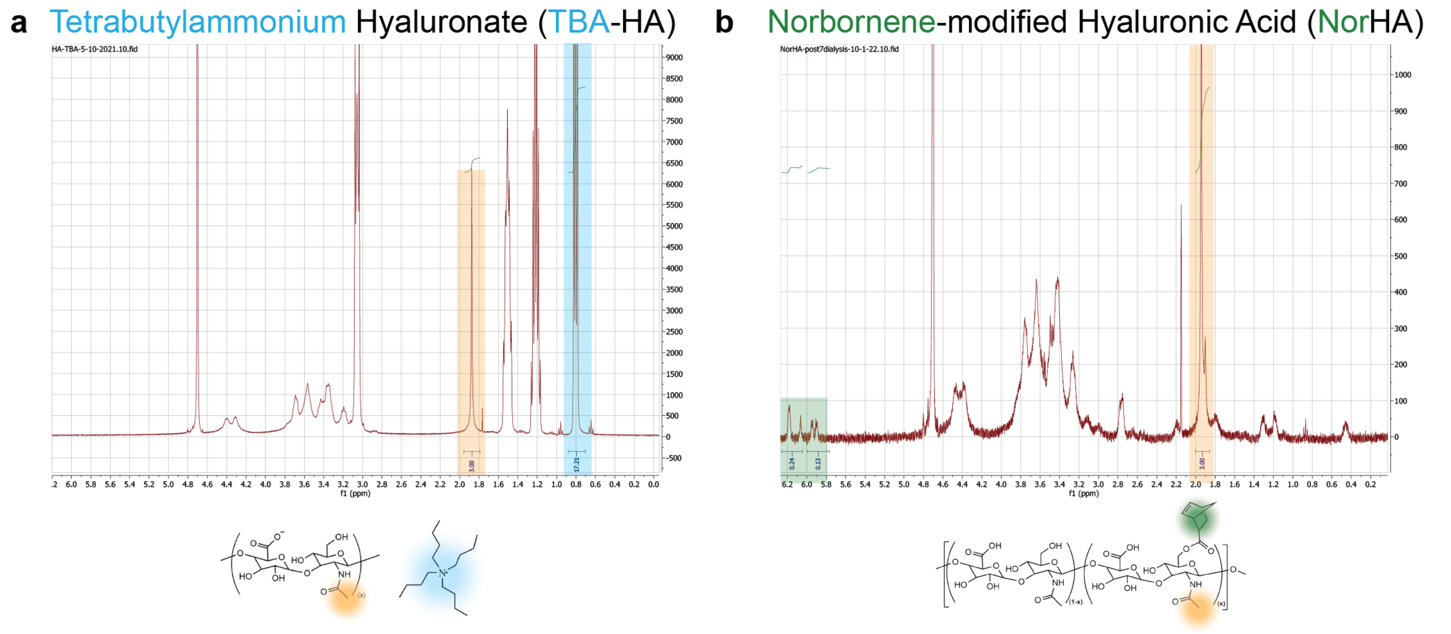


**Supplemental Figure 10 | $^{1}$H NMR spectra of synthesized TBA-HA and NorHA polymers in $D_2O$. a**, Representative NMR spectra of hyaluronic acid modified with tetrabutylammonium salt. Modification with tetrabutylammonium salt is determined by the integration of the TBA-methyl (blue) peak (12H, 0.6-0.9 ppm) relative to methyl group (orange) on HA (3H, 1.7-2.0 ppm). **b**, Representative NMR spectra of norbornene modified hyaluronic acid. Modification with norbornenes is determined by the integration of the vinyl (green) peaks (2H, 6.2-5.8 ppm) relative to methyl group (orange) on HA (3H, 1.7-2.0 ppm).

**Supplemental Table 1. qPCR gene of interest primer forward and reverse primers.**

| Number | Gene of Interest | Forward/Reverse | Sequence | Optical Density (OD) |
|---|---|---|---|---|
| 1 | COL2A1 | Forward | ACGCTCAAGTCCCTCAACAA | 5.37 |
| 2 | | Reverse | ACCTTCATGGCGTCCAAAGT | 6.57 |
| 3 | COL1A1 | Forward | GGGCAAGACAGTGATTGAATACA | 8.09 |
| 4 | | Reverse | GGATGGAGGGAGTTTACAGGAA | 7.20 |
| 5 | COL10A1 | Forward | AACGGGCAACAGCACTATGA | 5.13 |
| 6 | | Reverse | ACATGACAGGGGTGCCATTT | 5.20 |
| 7 | ACAN | Forward | AACTTAGCGGTGCCCATTCT | 6.31 |
| 8 | | Reverse | TGCAGAGGAGTCCCCACTTA | 6.00 |
| 9 | SOX9 | Forward | CACATCTCTCCCAACGCCAT | 6.48 |
| 10 | | Reverse | GTTGGTGGACCCTGGGATTG | 7.53 |
| 11 | GAPDH | Forward | GGTCACCAGGGCTGCTTTTA | 5.30 |
| 12 | | Reverse | TTGACTGTGCCGTGGAACTT | 5.96 |

### Supplemental Table 2. Statistical analysis report

| Figure | Panel | Statistical Test | Comparison | P-value | q/t value | Degrees of freedom |
|---|---|---|---|---|---|---|
| 2 | e: spheroids | | 20:80 vs. 35:65 | 0.6104 | 2.037 | 20 |
| | | | 20:80 vs. 50:50 | <0.0001 | 14.05 | 20 |
| | | | 20:80 vs. 65:35 | <0.0001 | 34.73 | 20 |
| | | ANOVA | 20:80 vs. 80:20 | <0.0001 | 35.19 | 20 |
| | | Tukey Multiple | 35:65 vs. 50:50 | <0.0001 | 12.01 | 20 |
| | | Comparisons | 35:65 vs. 65:35 | <0.0001 | 32.7 | 20 |
| | | Test | 35:65 vs. 80:20 | <0.0001 | 33.16 | 20 |
| | | | 50:50 vs. 65:35 | <0.0001 | 20.68 | 20 |
| | | | 50:50 vs. 80:20 | <0.0001 | 21.14 | 20 |
| | | | 65:35 vs. 80:20 | 0.9973 | 0.4614 | 20 |
| | e: microgels | | 20:80 vs. 35:65 | 0.9999 | 0.2167 | 20 |
| | | | 20:80 vs. 50:50 | <0.0001 | 18.9 | 20 |
| | | | 20:80 vs. 65:35 | <0.0001 | 29.12 | 20 |
| | | ANOVA | 20:80 vs. 80:20 | <0.0001 | 31.06 | 20 |
| | | Tukey Multiple | 35:65 vs. 50:50 | <0.0001 | 18.68 | 20 |
| | | Comparisons | 35:65 vs. 65:35 | <0.0001 | 28.91 | 20 |
| | | Test | 35:65 vs. 80:20 | <0.0001 | 30.84 | 20 |
| | | | 50:50 vs. 65:35 | <0.0001 | 10.23 | 20 |
| | | | 50:50 vs. 80:20 | <0.0001 | 12.16 | 20 |
| | | | 65:35 vs. 80:20 | 0.6545 | 1.933 | 20 |
| 3 | d | | Microgels vs. Spheroids | <0.0001 | 24.05 | 10 |
| | | | Microgels vs. 20:80 | 0.4687 | 2.421 | 10 |
| | | | Microgels vs. 35:65 | 0.4611 | 2.442 | 10 |
| | | ANOVA | Microgels vs. 50:50 | 0.8132 | 1.526 | 10 |
| | | Tukey Multiple | Spheroids vs. 20:80 | <0.0001 | 21.63 | 10 |
| | | Comparisons | Spheroids vs. 35:65 | <0.0001 | 26.49 | 10 |
| | | Test | Spheroids vs. 50:50 | <0.0001 | 25.58 | 10 |
| | | | 20:80 vs. 35:65 | 0.0397 | 4.863 | 10 |
| | | | 20:80 vs. 50:50 | 0.1079 | 3.947 | 10 |
| | | | 35:65 vs. 50:50 | 0.9632 | 0.9161 | 10 |
| | e | | Microgels vs. Spheroids | 0.9975 | 0.4458 | 10 |
| | | | Microgels vs. 20:80 | 0.209 | 3.311 | 10 |
| | | | Microgels vs. 35:65 | 0.001 | 8.398 | 10 |
| | | ANOVA | Microgels vs. 50:50 | 0.0003 | 9.959 | 10 |
| | | Tukey Multiple | Spheroids vs. 20:80 | 0.132 | 3.757 | 10 |
| | | Comparisons | Spheroids vs. 35:65 | 0.0007 | 8.844 | 10 |
| | | Test | Spheroids vs. 50:50 | 0.0002 | 10.41 | 10 |
| | | | 20:80 vs. 35:65 | 0.0311 | 5.087 | 10 |
| | | | 20:80 vs. 50:50 | 0.0059 | 6.648 | 10 |
| | | | 35:65 vs. 50:50 | 0.8011 | 1.561 | 10 |
| 4 | c | Two-way ANOVA | 20:80:microgels vs. 20:80:spheroids | <0.0001 | 25.55 | 24 |
| | | Tukey Multiple | 20:80:microgels vs. 35:65:microgels | 0.0086 | 4.995 | 24 |
| | | Comparisons | 20:80:spheroids vs. 35:65:spheroids | 0.0086 | 4.995 | 24 |
| | | Test | 35:65:microgels vs. 35:65:spheroids | <0.0001 | 12.78 | 24 |
| | d:pore area | Mann Whitney test | 20:80 vs 35:65 | 0.0175 | N/A | N/A |
| | e: % area fraction | | 20:80 vs 35:65 | 0.3357 | N/A | N/A |
| 5 | b | | Day 1:20:80 vs. Day 1:35:65 | 0.8514 | 1.648 | 66 |
| | | | Day 1:20:80 vs. Day 28:20:80 | >0.9999 | 0.2189 | 66 |
| | | | Day 1:20:80 vs. Day 28:35:65 | >0.9999 | 0.2811 | 66 |
| | | | Day 1:20:80 vs. Day 56:20:80 | 0.8253 | 1.726 | 66 |
| | | | Day 1:20:80 vs. Day 56:35:65 | 0.9365 | 1.319 | 66 |
| | | Two-way ANOVA | Day 1:35:65 vs. Day 28:20:80 | 0.7728 | 1.867 | 66 |
| | | Tukey Multiple | Day 1:35:65 vs. Day 28:35:65 | 0.7478 | 1.929 | 66 |
| | | Comparisons | Day 1:35:65 vs. Day 56:20:80 | >0.9999 | 0.07775 | 66 |
| | | Test | Day 1:35:65 vs. Day 56:35:65 | >0.9999 | 0.329 | 66 |
| | | | Day 28:20:80 vs. Day 28:35:65 | >0.9999 | 0.0622 | 66 |
| | | | Day 28:20:80 vs. Day 56:20:80 | 0.7414 | 1.945 | 66 |

| | | | | | |
|---|---|---|---|---|---|
| | | Day 28:20:80 vs. Day 56:35:65 | 0.8846 | 1.538 | 66 |
| | | Day 28:35:65 vs. Day 56:20:80 | 0.7153 | 2.007 | 66 |
| | | Day 28:35:65 vs. Day 56:35:65 | 0.8664 | 1.601 | 66 |
| | | Day 56:20:80 vs. Day 56:35:65 | 0.9997 | 0.4067 | 66 |
| c i. dsDNA | Two-way ANOVA Tukey Multiple Comparisons Test | Day 1:20:80 vs. Day 1:35:65 | 0.402 | 2.715 | 52 |
| | | Day 1:20:80 vs. Day 28:20:80 | 0.808 | 1.773 | 52 |
| | | Day 1:20:80 vs. Day 28:35:65 | 0.0028 | 5.624 | 52 |
| | | Day 1:20:80 vs. Day 56:20:80 | 0.9846 | 0.9456 | 52 |
| | | Day 1:20:80 vs. Day 56:35:65 | 0.1309 | 3.595 | 52 |
| | | Day 1:35:65 vs. Day 28:20:80 | 0.9792 | 1.012 | 52 |
| | | Day 1:35:65 vs. Day 28:35:65 | 0.3255 | 2.909 | 52 |
| | | Day 1:35:65 vs. Day 56:20:80 | 0.1062 | 3.731 | 52 |
| | | Day 1:35:65 vs. Day 56:35:65 | 0.9948 | 0.7469 | 52 |
| | | Day 28:20:80 vs. Day 28:35:65 | 0.069 | 3.997 | 52 |
| | | Day 28:20:80 vs. Day 56:20:80 | 0.3701 | 2.793 | 52 |
| | | Day 28:20:80 vs. Day 56:35:65 | 0.7858 | 1.833 | 52 |
| | | Day 28:35:65 vs. Day 56:20:80 | 0.0002 | 6.716 | 52 |
| | | Day 28:35:65 vs. Day 56:35:65 | 0.5835 | 2.304 | 52 |
| | | Day 56:20:80 vs. Day 56:35:65 | 0.0195 | 4.692 | 52 |
| c ii. sGAG | Two-way ANOVA Tukey Multiple Comparisons Test | Day 1:20:80 vs. Day 1:35:65 | 0.9897 | 0.8651 | 52 |
| | | Day 1:20:80 vs. Day 28:20:80 | 0.0035 | 5.524 | 52 |
| | | Day 1:20:80 vs. Day 28:35:65 | <0.0001 | 9.067 | 52 |
| | | Day 1:20:80 vs. Day 56:20:80 | 0.0001 | 6.976 | 52 |
| | | Day 1:20:80 vs. Day 56:35:65 | <0.0001 | 15 | 52 |
| | | Day 1:35:65 vs. Day 28:20:80 | 0.0005 | 6.411 | 52 |
| | | Day 1:35:65 vs. Day 28:35:65 | <0.0001 | 9.932 | 52 |
| | | Day 1:35:65 vs. Day 56:20:80 | <0.0001 | 7.864 | 52 |
| | | Day 1:35:65 vs. Day 56:35:65 | <0.0001 | 15.91 | 52 |
| | | Day 28:20:80 vs. Day 28:35:65 | 0.0985 | 3.779 | 52 |
| | | Day 28:20:80 vs. Day 56:20:80 | 0.8965 | 1.493 | 52 |
| | | Day 28:20:80 vs. Day 56:35:65 | <0.0001 | 9.622 | 52 |
| | | Day 28:35:65 vs. Day 56:20:80 | 0.5736 | 2.326 | 52 |
| | | Day 28:35:65 vs. Day 56:35:65 | 0.0038 | 5.491 | 52 |
| | | Day 56:20:80 vs. Day 56:35:65 | <0.0001 | 8.095 | 52 |
| c iii. Collagen | Two-way ANOVA Tukey Multiple Comparisons Test | Day 1:20:80 vs. Day 1:35:65 | 0.9532 | 1.208 | 21 |
| | | Day 1:20:80 vs. Day 28:20:80 | 0.0001 | 8.124 | 21 |
| | | Day 1:20:80 vs. Day 28:35:65 | <0.0001 | 8.372 | 21 |
| | | Day 1:20:80 vs. Day 56:20:80 | <0.0001 | 12 | 21 |
| | | Day 1:20:80 vs. Day 56:35:65 | <0.0001 | 15.48 | 21 |
| | | Day 1:35:65 vs. Day 28:20:80 | 0.0011 | 6.85 | 21 |
| | | Day 1:35:65 vs. Day 28:35:65 | 0.0006 | 7.164 | 21 |
| | | Day 1:35:65 vs. Day 56:20:80 | <0.0001 | 10.73 | 21 |
| | | Day 1:35:65 vs. Day 56:35:65 | <0.0001 | 14.21 | 21 |
| | | Day 28:20:80 vs. Day 28:35:65 | 0.9958 | 0.701 | 21 |
| | | Day 28:20:80 vs. Day 56:20:80 | 0.0785 | 4.111 | 21 |
| | | Day 28:20:80 vs. Day 56:35:65 | 0.0002 | 7.802 | 21 |
| | | Day 28:35:65 vs. Day 56:20:80 | 0.2598 | 3.175 | 21 |
| | | Day 28:35:65 vs. Day 56:35:65 | 0.0015 | 6.655 | 21 |
| | | Day 56:20:80 vs. Day 56:35:65 | 0.1385 | 3.691 | 21 |
| d | Two-way ANOVA Tukey Multiple Comparisons Test | Day 28:20:80 vs. Day 28:35:65 | 0.9546 | 0.7294 | 28 |
| | | Day 28:20:80 vs. Day 56:20:80 | <0.0001 | 8.078 | 28 |
| | | Day 28:20:80 vs. Day 56:35:65 | <0.0001 | 12.68 | 28 |
| | | Day 28:35:65 vs. Day 56:20:80 | <0.0001 | 8.807 | 28 |
| | | Day 28:35:65 vs. Day 56:35:65 | <0.0001 | 13.41 | 28 |
| | | Day 56:20:80 vs. Day 56:35:65 | 0.0148 | 4.602 | 28 |
| | Two-tailed T-test | Day 1 20:80 vs Day 1 35:65 | 0.0003 | 6.046 | 8 |
| f | Mann Whitney test | sGAG area fraction 20:80 vs 35:65 | 0.0286 | N/A | N/A |
| | | Col 2 area fraction 20:80 vs 35:65 | 0.0286 | N/A | N/A |
| | | sGAG integrated density 20:80 vs 35:65 | 0.0286 | N/A | N/A |
| | | Col 2 integrated density 20:80 vs 35:65 | 0.0286 | N/A | N/A |

| | | | | | | |
|---|---|---|---|---|---|---|
| 6 | b | Two-tailed T-test | MSC Only vs Co-culture | 0.1246 | 1.6077 | 19 |
| | c | Two-way ANOVA Tukey Multiple Comparisons Test | Day 1:MSC Only vs. Day 1: Co-Culture | >0.9999 | 0.1462 | 23 |
| | | | Day 1:MSC Only vs. Day 28:MSC Only | 0.513 | 2.479 | 23 |
| | | | Day 1:MSC Onlyvs. Day 28: Co-Culture | <0.0001 | 10.73 | 23 |
| | | | Day 1:MSC Only vs. Day 56:MSC Only | 0.285 | 3.081 | 23 |
| | | | Day 1:MSC Only vs. Day 56: Co-Culture | <0.0001 | 12.1 | 23 |
| | | | Day 1: Co-Culture vs. Day 28:MSC Only | 0.5667 | 2.355 | 23 |
| | | | Day 1: Co-Culture vs. Day 28: Co-Culture | <0.0001 | 10.85 | 23 |
| | | | Day 1: Co-Culture vs. Day 56:MSC Only | 0.3263 | 2.957 | 23 |
| | | | Day 1: Co-Culture vs. Day 56: Co-Culture | <0.0001 | 12.22 | 23 |
| | | | Day 28:MSC Onlyvs. Day 28:Co-Culture | <0.0001 | 11.73 | 23 |
| | | | Day 28:MSC Only vs. Day 56:MSC Only | 0.9989 | 0.5335 | 23 |
| | | | Day 28:MSC Only vs. Day 56: Co-Culture | <0.0001 | 12.92 | 23 |
| | | | Day 28: Co-Culture vs. Day 56:MSC Only | <0.0001 | 12.23 | 23 |
| | | | Day 28: Co-Culture vs. Day 56: Co-Culture | >0.9999 | 0.23 | 23 |
| | | | Day 56:MSC Only vs. Day 56: Co-Culture | <0.0001 | 13.45 | 23 |
| | d i. dsDNA | Two-way ANOVA Tukey Multiple Comparisons Test | Day 1:MSC Only vs. Day 1: Co-Culture | 0.9812 | 0.9679 | 16 |
| | | | Day 1:MSC Only vs. Day 28:MSC Only | 0.9928 | 0.7811 | 16 |
| | | | Day 1:MSC Onlyvs. Day 28: Co-Culture | 0.0001 | 9.058 | 16 |
| | | | Day 1:MSC Only vs. Day 56:MSC Only | >0.9999 | 0.3047 | 16 |
| | | | Day 1:MSC Only vs. Day 56: Co-Culture | 0.107 | 3.972 | 16 |
| | | | Day 1: Co-Culture vs. Day 28:MSC Only | 0.7619 | 1.889 | 16 |
| | | | Day 1: Co-Culture vs. Day 28: Co-Culture | <0.0001 | 10.65 | 16 |
| | | | Day 1: Co-Culture vs. Day 56:MSC Only | 0.9203 | 1.375 | 16 |
| | | | Day 1: Co-Culture vs. Day 56: Co-Culture | 0.0172 | 5.336 | 16 |
| | | | Day 28:MSC Onlyvs. Day 28:Co-Culture | 0.0001 | 8.902 | 16 |
| | | | Day 28:MSC Only vs. Day 56:MSC Only | 0.999 | 0.5146 | 16 |
| | | | Day 28:MSC Only vs. Day 56: Co-Culture | 0.2012 | 3.447 | 16 |
| | | | Day 28: Co-Culture vs. Day 56:MSC Only | <0.0001 | 9.378 | 16 |
| | | | Day 28: Co-Culture vs. Day 56: Co-Culture | 0.0102 | 5.711 | 16 |
| | | | Day 56:MSC Only vs. Day 56: Co-Culture | 0.1085 | 3.962 | 16 |
| | d ii. sGAG | Two-way ANOVA Tukey Multiple Comparisons Test | Day 1:MSC Only vs. Day 1: Co-Culture | 0.3708 | 2.865 | 16 |
| | | | Day 1:MSC Only vs. Day 28:MSC Only | <0.0001 | 10.34 | 16 |
| | | | Day 1:MSC Onlyvs. Day 28: Co-Culture | <0.0001 | 13.35 | 16 |
| | | | Day 1:MSC Only vs. Day 56:MSC Only | <0.0001 | 12.03 | 16 |
| | | | Day 1:MSC Only vs. Day 56: Co-Culture | <0.0001 | 16.54 | 16 |
| | | | Day 1: Co-Culture vs. Day 28:MSC Only | 0.0004 | 8.07 | 16 |
| | | | Day 1: Co-Culture vs. Day 28: Co-Culture | <0.0001 | 11.41 | 16 |
| | | | Day 1: Co-Culture vs. Day 56:MSC Only | <0.0001 | 9.901 | 16 |
| | | | Day 1: Co-Culture vs. Day 56: Co-Culture | <0.0001 | 14.77 | 16 |
| | | | Day 28:MSC Onlyvs. Day 28:Co-Culture | 0.1114 | 3.94 | 16 |
| | | | Day 28:MSC Only vs. Day 56:MSC Only | 0.7839 | 1.83 | 16 |
| | | | Day 28:MSC Only vs. Day 56: Co-Culture | 0.0025 | 6.703 | 16 |
| | | | Day 28: Co-Culture vs. Day 56:MSC Only | 0.6173 | 2.245 | 16 |
| | | | Day 28: Co-Culture vs. Day 56: Co-Culture | 0.6087 | 2.266 | 16 |
| | | | Day 56:MSC Only vs. Day 56: Co-Culture | 0.0326 | 4.873 | 16 |
| | d iii. Collagen | Two-way ANOVA Tukey Multiple Comparisons Test | Day 1:MSC Only vs. Day 1: Co-Culture | 0.0545 | 4.492 | 16 |
| | | | Day 1:MSC Only vs. Day 28:MSC Only | 0.0015 | 7.084 | 16 |
| | | | Day 1:MSC Onlyvs. Day 28: Co-Culture | 0.0017 | 6.986 | 16 |
| | | | Day 1:MSC Only vs. Day 56:MSC Only | <0.0001 | 12.44 | 16 |
| | | | Day 1:MSC Only vs. Day 56: Co-Culture | <0.0001 | 10.07 | 16 |
| | | | Day 1: Co-Culture vs. Day 28:MSC Only | 0.3944 | 2.799 | 16 |
| | | | Day 1: Co-Culture vs. Day 28: Co-Culture | 0.3328 | 2.976 | 16 |
| | | | Day 1: Co-Culture vs. Day 56:MSC Only | 0.0002 | 8.58 | 16 |
| | | | Day 1: Co-Culture vs. Day 56: Co-Culture | 0.0065 | 6.025 | 16 |
| | | | Day 28:MSC Onlyvs. Day 28:Co-Culture | 0.9998 | 0.3846 | 16 |
| | | | Day 28:MSC Only vs. Day 56:MSC Only | 0.0092 | 5.781 | 16 |
| | | | Day 28:MSC Only vs. Day 56: Co-Culture | 0.2572 | 3.225 | 16 |
| | | | Day 28: Co-Culture vs. Day 56:MSC Only | 0.0286 | 4.967 | 16 |
| | | | Day 28: Co-Culture vs. Day 56: Co-Culture | 0.4702 | 2.601 | 16 |
| | | | Day 56:MSC Only vs. Day 56: Co-Culture | 0.4885 | 2.555 | 16 |

| | | | | | |
|---|---|---|---|---|---|
| e | | Day 28:MSC Only vs. Day 28:Co-Culture | 0.278 | 2.656 | 15 |
| | Two-way ANOVA | Day 28:MSC Only vs. Day 56:MSC Only | <0.0001 | 11.11 | 15 |
| | Tukey Multiple | Day 28:MSC Only vs. Day 56:Co-Culture | <0.0001 | 15.29 | 15 |
| | Comparisons | Day 28:Co-Culture vs. Day 56:MSC Only | <0.0001 | 8.97 | 15 |
| | Test | Day 28:Co-Culture vs. Day 56:Co-Culture | <0.0001 | 13.4 | 15 |
| | | Day 56:MSC Only vs. Day 56:Co-Culture | 0.0312 | 4.427 | 15 |
| | Two-tailed T-test | Day 1: MSC Only vs. Day 1: Co-Culture | 0.2773 | 1.257 | 4 |
| g | | sGAG area fraction MSC Only vs Co-Culture | 0.0286 N/A | N/A | |
| | Mann Whitney | Col 2 area fraction MSC Only vs. Co-Culture | 0.0286 N/A | N/A | |
| | test | sGAG integrated density MSC Only vs Co-Culture | 0.4857 N/A | N/A | |
| | | Col 2 integrated density MSC Only vs. Co-Culture | 0.1143 N/A | N/A | |
| Supplemental Figures | | | | | |
| 1 a | ANOVA | 500 vs. 1000 | <0.0001 | 8.043 | 147 |
| | Tukey Multiple | 500 vs. 2000 | <0.0001 | 31.64 | 147 |
| | Comparisons | 1000 vs. 2000 | <0.0001 | 23.6 | 147 |
| c | ANOVA | 200 vs. 280 | <0.0001 | 12.97 | 284 |
| | Tukey Multiple | 200 vs. 350 | <0.0001 | 19.98 | 284 |
| | Comparisons | 280 vs. 350 | <0.0001 | 10.63 | 284 |
| 2 | | Day 1:500 vs. Day 1:1000 | 0.1486 | 3.468 | 102 |
| | | Day 1:500 vs. Day 1:2000 | <0.0001 | 15.87 | 102 |
| | | Day 1:500 vs. Day 7:500 | 0.9148 | 1.424 | 102 |
| | | Day 1:500 vs. Day 7:1000 | 0.1009 | 3.711 | 102 |
| | | Day 1:500 vs. Day 7:2000 | <0.0001 | 9.081 | 102 |
| | | Day 1:1000 vs. Day 1:2000 | <0.0001 | 12.4 | 102 |
| | Two-way ANOVA | Day 1:1000 vs. Day 7:500 | 0.6992 | 2.044 | 102 |
| | Tukey Multiple | Day 1:1000 vs. Day 7:1000 | >0.9999 | 0.2427 | 102 |
| | Comparisons | Day 1:1000 vs. Day 7:2000 | 0.0018 | 5.613 | 102 |
| | Test | Day 1:2000 vs. Day 7:500 | <0.0001 | 14.44 | 102 |
| | | Day 1:2000 vs. Day 7:1000 | <0.0001 | 12.16 | 102 |
| | | Day 1:2000 vs. Day 7:2000 | <0.0001 | 6.786 | 102 |
| | | Day 7:500 vs. Day 7:1000 | 0.5893 | 2.287 | 102 |
| | | Day 7:500 vs. Day 7:2000 | <0.0001 | 7.658 | 102 |
| | | Day 7:1000 vs. Day 7:2000 | 0.0033 | 5.371 | 102 |
| 3 a | | Day 1:donor 1 vs. Day 1:donor 2 | 0.9992 | 1.52 | 466 |
| | | Day 1:donor 1 vs. Day 1:donor 3 | 0.9951 | 1.793 | 466 |
| | | Day 1:donor 1 vs. Day 7:donor 1 | 0.0001 | 6.953 | 466 |
| | | Day 1:donor 1 vs. Day 7:donor 2 | <0.0001 | 11.47 | 466 |
| | | Day 1:donor 1 vs. Day 7:donor 3 | >0.9999 | 0.6327 | 466 |
| | | Day 1:donor 1 vs. Day 14:donor 1 | <0.0001 | 7.498 | 466 |
| | | Day 1:donor 1 vs. Day 14:donor 2 | 0.0001 | 6.989 | 466 |
| | | Day 1:donor 1 vs. Day 14:donor 3 | 0.0013 | 6.218 | 466 |
| | | Day 1:donor 1 vs. Day 21:donor 1 | <0.0001 | 7.963 | 466 |
| | | Day 1:donor 1 vs. Day 21:donor 2 | 0.0003 | 6.716 | 466 |
| | | Day 1:donor 1 vs. Day 21:donor 3 | <0.0001 | 7.147 | 466 |
| | | Day 1:donor 1 vs. Day 28:donor 1 | <0.0001 | 9.582 | 466 |
| | | Day 1:donor 1 vs. Day 28:donor 2 | <0.0001 | 8.355 | 466 |
| | | Day 1:donor 1 vs. Day 28:donor 3 | <0.0001 | 12.52 | 466 |
| | | Day 1:donor 2 vs. Day 1:donor 3 | 0.559 | 3.313 | 466 |
| | | Day 1:donor 2 vs. Day 7:donor 1 | 0.0156 | 5.317 | 466 |
| | | Day 1:donor 2 vs. Day 7:donor 2 | <0.0001 | 9.834 | 466 |
| | | Day 1:donor 2 vs. Day 7:donor 3 | 0.9572 | 2.269 | 466 |
| | | Day 1:donor 2 vs. Day 14:donor 1 | 0.0032 | 5.906 | 466 |
| | | Day 1:donor 2 vs. Day 14:donor 2 | 0.0064 | 5.658 | 466 |
| | | Day 1:donor 2 vs. Day 14:donor 3 | 0.0433 | 4.888 | 466 |
| | | Day 1:donor 2 vs. Day 21:donor 1 | 0.0003 | 6.711 | 466 |
| | | Day 1:donor 2 vs. Day 21:donor 2 | 0.0107 | 5.464 | 466 |
| | | Day 1:donor 2 vs. Day 21:donor 3 | 0.0033 | 5.895 | 466 |
| | | Day 1:donor 2 vs. Day 28:donor 1 | <0.0001 | 8.251 | 466 |
| | | Day 1:donor 2 vs. Day 28:donor 2 | <0.0001 | 7.024 | 466 |

Two-way ANOVA Tukey Multiple Comparisons Test

| | | | |
|---|---|---|---|
| Day 1:donor 2 vs. Day 28:donor 3 | <0.0001 | 11.19 | 466 |
| Day 1:donor 3 vs. Day 7:donor 1 | <0.0001 | 8.883 | 466 |
| Day 1:donor 3 vs. Day 7:donor 2 | <0.0001 | 13.4 | 466 |
| Day 1:donor 3 vs. Day 7:donor 3 | 0.9999 | 1.298 | 466 |
| Day 1:donor 3 vs. Day 14:donor 1 | <0.0001 | 9.377 | 466 |
| Day 1:donor 3 vs. Day 14:donor 2 | <0.0001 | 8.559 | 466 |
| Day 1:donor 3 vs. Day 14:donor 3 | <0.0001 | 7.789 | 466 |
| Day 1:donor 3 vs. Day 21:donor 1 | <0.0001 | 9.44 | 466 |
| Day 1:donor 3 vs. Day 21:donor 2 | <0.0001 | 8.193 | 466 |
| Day 1:donor 3 vs. Day 21:donor 3 | <0.0001 | 8.624 | 466 |
| Day 1:donor 3 vs. Day 28:donor 1 | <0.0001 | 11.15 | 466 |
| Day 1:donor 3 vs. Day 28:donor 2 | <0.0001 | 9.925 | 466 |
| Day 1:donor 3 vs. Day 28:donor 3 | <0.0001 | 14.09 | 466 |
| Day 7:donor 1 vs. Day 7:donor 2 | 0.0397 | 4.926 | 466 |
| Day 7:donor 1 vs. Day 7:donor 3 | <0.0001 | 8.272 | 466 |
| Day 7:donor 1 vs. Day 14:donor 1 | >0.9999 | 0.7947 | 466 |
| Day 7:donor 1 vs. Day 14:donor 2 | 0.9996 | 1.41 | 466 |
| Day 7:donor 1 vs. Day 14:donor 3 | >0.9999 | 0.5959 | 466 |
| Day 7:donor 1 vs. Day 21:donor 1 | 0.8192 | 2.775 | 466 |
| Day 7:donor 1 vs. Day 21:donor 2 | 0.9994 | 1.466 | 466 |
| Day 7:donor 1 vs. Day 21:donor 3 | 0.9904 | 1.919 | 466 |
| Day 7:donor 1 vs. Day 28:donor 1 | 0.184 | 4.152 | 466 |
| Day 7:donor 1 vs. Day 28:donor 2 | 0.7865 | 2.855 | 466 |
| Day 7:donor 1 vs. Day 28:donor 3 | <0.0001 | 7.257 | 466 |
| Day 7:donor 2 vs. Day 7:donor 3 | <0.0001 | 13.2 | 466 |
| Day 7:donor 2 vs. Day 14:donor 1 | 0.2448 | 3.975 | 466 |
| Day 7:donor 2 vs. Day 14:donor 2 | 0.9165 | 2.473 | 466 |
| Day 7:donor 2 vs. Day 14:donor 3 | 0.5725 | 3.288 | 466 |
| Day 7:donor 2 vs. Day 21:donor 1 | >0.9999 | 0.8539 | 466 |
| Day 7:donor 2 vs. Day 21:donor 2 | 0.9713 | 2.163 | 466 |
| Day 7:donor 2 vs. Day 21:donor 3 | 0.997 | 1.711 | 466 |
| Day 7:donor 2 vs. Day 28:donor 1 | >0.9999 | 0.268 | 466 |
| Day 7:donor 2 vs. Day 28:donor 2 | >0.9999 | 1.029 | 466 |
| Day 7:donor 2 vs. Day 28:donor 3 | 0.5271 | 3.373 | 466 |
| Day 7:donor 3 vs. Day 14:donor 1 | <0.0001 | 8.804 | 466 |
| Day 7:donor 3 vs. Day 14:donor 2 | <0.0001 | 7.932 | 466 |
| Day 7:donor 3 vs. Day 14:donor 3 | <0.0001 | 7.118 | 466 |
| Day 7:donor 3 vs. Day 21:donor 1 | <0.0001 | 8.87 | 466 |
| Day 7:donor 3 vs. Day 21:donor 2 | <0.0001 | 7.561 | 466 |
| Day 7:donor 3 vs. Day 21:donor 3 | <0.0001 | 8.013 | 466 |
| Day 7:donor 3 vs. Day 28:donor 1 | <0.0001 | 10.67 | 466 |
| Day 7:donor 3 vs. Day 28:donor 2 | <0.0001 | 9.376 | 466 |
| Day 7:donor 3 vs. Day 28:donor 3 | <0.0001 | 13.78 | 466 |
| Day 14:donor 1 vs. Day 14:donor 2 | >0.9999 | 0.7479 | 466 |
| Day 14:donor 1 vs. Day 14:donor 3 | >0.9999 | 0.05015 | 466 |
| Day 14:donor 1 vs. Day 21:donor 1 | 0.9747 | 2.132 | 466 |
| Day 14:donor 1 vs. Day 21:donor 2 | >0.9999 | 0.8461 | 466 |
| Day 14:donor 1 vs. Day 21:donor 3 | 0.9999 | 1.291 | 466 |
| Day 14:donor 1 vs. Day 28:donor 1 | 0.495 | 3.434 | 466 |
| Day 14:donor 1 vs. Day 28:donor 2 | 0.9713 | 2.163 | 466 |
| Day 14:donor 1 vs. Day 28:donor 3 | 0.0006 | 6.477 | 466 |
| Day 14:donor 2 vs. Day 14:donor 3 | >0.9999 | 0.6937 | 466 |
| Day 14:donor 2 vs. Day 21:donor 1 | 0.9999 | 1.264 | 466 |
| Day 14:donor 2 vs. Day 21:donor 2 | >0.9999 | 0.1283 | 466 |
| Day 14:donor 2 vs. Day 21:donor 3 | >0.9999 | 0.5207 | 466 |
| Day 14:donor 2 vs. Day 28:donor 1 | 0.9461 | 2.335 | 466 |
| Day 14:donor 2 vs. Day 28:donor 2 | >0.9999 | 1.23 | 466 |
| Day 14:donor 2 vs. Day 28:donor 3 | 0.0351 | 4.98 | 466 |
| Day 14:donor 3 vs. Day 21:donor 1 | 0.9902 | 1.923 | 466 |
| Day 14:donor 3 vs. Day 21:donor 2 | >0.9999 | 0.7882 | 466 |
| Day 14:donor 3 vs. Day 21:donor 3 | >0.9999 | 1.181 | 466 |
| Day 14:donor 3 vs. Day 28:donor 1 | 0.7062 | 3.029 | 466 |
| Day 14:donor 3 vs. Day 28:donor 2 | 0.9902 | 1.924 | 466 |
| Day 14:donor 3 vs. Day 28:donor 3 | 0.0061 | 5.674 | 466 |
| Day 21:donor 1 vs. Day 21:donor 2 | >0.9999 | 1.085 | 466 |
| Day 21:donor 1 vs. Day 21:donor 3 | >0.9999 | 0.7098 | 466 |

| | | | | | |
|---|---|---|---|---|---|
| | | Day 21:donor 1 vs. Day 28:donor 1 | >0.9999 | 0.9574 | 466 |
| | | Day 21:donor 1 vs. Day 28:donor 2 | >0.9999 | 0.09334 | 466 |
| | | Day 21:donor 1 vs. Day 28:donor 3 | 0.4746 | 3.473 | 466 |
| | | Day 21:donor 2 vs. Day 21:donor 3 | >0.9999 | 0.375 | 466 |
| | | Day 21:donor 2 vs. Day 28:donor 1 | 0.9786 | 2.093 | 466 |
| | | Day 21:donor 2 vs. Day 28:donor 2 | >0.9999 | 1.042 | 466 |
| | | Day 21:donor 2 vs. Day 28:donor 3 | 0.0785 | 4.608 | 466 |
| | | Day 21:donor 3 vs. Day 28:donor 1 | 0.9972 | 1.7 | 466 |
| | | Day 21:donor 3 vs. Day 28:donor 2 | >0.9999 | 0.6495 | 466 |
| | | Day 21:donor 3 vs. Day 28:donor 3 | 0.1648 | 4.216 | 466 |
| | | Day 28:donor 1 vs. Day 28:donor 2 | >0.9999 | 1.105 | 466 |
| | | Day 28:donor 1 vs. Day 28:donor 3 | 0.8667 | 2.645 | 466 |
| | | Day 28:donor 2 vs. Day 28:donor 3 | 0.3387 | 3.75 | 466 |
| b | | Day 1:Donor 2 vs. Day 1:Donor 3 | 0.9985 | 0.5826 | 174 |
| | | Day 1:Donor 2 vs. Day 1:Donor 4 | 0.9573 | 1.204 | 174 |
| | | Day 1:Donor 2 vs. Day 28:Donor 2 | <0.0001 | 8.242 | 174 |
| | | Day 1:Donor 2 vs. Day 28:Donor 3 | <0.0001 | 7.098 | 174 |
| | | Day 1:Donor 2 vs. Day 28:Donor 4 | <0.0001 | 11.56 | 174 |
| | | Day 1:Donor 3 vs. Day 1:Donor 4 | 0.8046 | 1.786 | 174 |
| | Two-way ANOVA | Day 1:Donor 3 vs. Day 28:Donor 2 | <0.0001 | 7.732 | 174 |
| | Tukey Multiple | Day 1:Donor 3 vs. Day 28:Donor 3 | <0.0001 | 6.587 | 174 |
| | Comparisons | Day 1:Donor 3 vs. Day 28:Donor 4 | <0.0001 | 11.05 | 174 |
| | Test | Day 1:Donor 4 vs. Day 28:Donor 2 | <0.0001 | 9.295 | 174 |
| | | Day 1:Donor 4 vs. Day 28:Donor 3 | <0.0001 | 8.151 | 174 |
| | | Day 1:Donor 4 vs. Day 28:Donor 4 | <0.0001 | 12.61 | 174 |
| | | Day 28:Donor 2 vs. Day 28:Donor 3 | 0.9782 | 1.03 | 174 |
| | | Day 28:Donor 2 vs. Day 28:Donor 4 | 0.2852 | 2.989 | 174 |
| | | Day 28:Donor 3 vs. Day 28:Donor 4 | 0.0557 | 4.019 | 174 |
| 4 c: microgels | | 2080:(-) vs. 2080:+ SD | <0.0001 | 13.61 | 16 |
| | Two-way ANOVA | 2080:(-) vs. 3565:(-) | 0.925 | 0.8733 | 16 |
| | Tukey Multiple | 2080:(-) vs. 3565:+ SD | <0.0001 | 10.42 | 16 |
| | Comparisons | 2080:+ SD vs. 3565:(-) | <0.0001 | 12.74 | 16 |
| | Test | 2080:+ SD vs. 3565:+ SD | 0.1509 | 3.188 | 16 |
| | | 3565:(-) vs. 3565:+ SD | <0.0001 | 9.551 | 16 |
| c: spheroids | | 2080:(-) vs. 2080:+ SD | 0.9977 | 0.2584 | 16 |
| | Two-way ANOVA | 2080:(-) vs. 3565:(-) | 0.7358 | 1.454 | 16 |
| | Tukey Multiple | 2080:(-) vs. 3565:+ SD | 0.0595 | 3.917 | 16 |
| | Comparisons | 2080:+ SD vs. 3565:(-) | 0.8321 | 1.196 | 16 |
| | Test | 2080:+ SD vs. 3565:+ SD | 0.0837 | 3.658 | 16 |
| | | 3565:(-) vs. 3565:+ SD | 0.3359 | 2.463 | 16 |
| d: microgels | | 20:80:(-) vs. 20:80:(+) | >0.9999 | 0.03651 | 16 |
| | Two-way ANOVA | 20:80:(-) vs. 35:65:(-) | >0.9999 | 0.08796 | 16 |
| | Tukey Multiple | 20:80:(-) vs. 35:65:(+) | 0.1465 | 3.213 | 16 |
| | Comparisons | 20:80:(+) vs. 35:65:(-) | 0.9997 | 0.1245 | 16 |
| | Test | 20:80:(+) vs. 35:65:(+) | 0.1401 | 3.25 | 16 |
| | | 35:65:(-) vs. 35:65:(+) | 0.1628 | 3.125 | 16 |
| d: spheorids | | 20:80:(-) vs. 20:80:(+) | 0.4065 | 2.262 | 16 |
| | Two-way ANOVA | 20:80:(-) vs. 35:65:(-) | 0.6277 | 1.716 | 16 |
| | Tukey Multiple | 20:80:(-) vs. 35:65:(+) | <0.0001 | 9.546 | 16 |
| | Comparisons | 20:80:(+) vs. 35:65:(-) | 0.9797 | 0.5464 | 16 |
| | Test | 20:80:(+) vs. 35:65:(+) | 0.0005 | 7.284 | 16 |
| | | 35:65:(-) vs. 35:65:(+) | 0.0002 | 7.83 | 16 |
| 5 a: G' | ANOVA | 500 vs. 15k | 0.0009 | 7.927 | 9 |
| | Tukey Multiple | 500 vs. vaccum | <0.0001 | 14.52 | 9 |
| | Comparisons | 15k vs. vaccum | 0.0031 | 6.588 | 9 |
| a: G" | ANOVA | 500 vs. 15k | 0.0494 | 3.959 | 9 |
| | Tukey Multiple | 500 vs. vaccum | 0.0003 | 9.308 | 9 |
| | Comparisons | 15k vs. vaccum | 0.0109 | 5.349 | 9 |

| | | | | | |
|---|---|---|---|---|---|
| b | ANOVA Tukey Multiple Comparisons Test | 0.5 vs. 1 min. | 0.0192 | 4.973 | 12 |
| | | 0.5 vs. 3 min. | 0.0009 | 7.509 | 12 |
| | | 0.5 vs. 5 min. | 0.0002 | 8.822 | 12 |
| | | 1 vs. 3 min. | 0.3229 | 2.536 | 12 |
| | | 1 vs. 5 min. | 0.0763 | 3.849 | 12 |
| | | 3 vs. 5 min. | 0.7905 | 1.313 | 12 |
| 7 a: SOX9 | Two-way ANOVA Tukey Multiple Comparisons Test | Day 1:20:80 vs. Day 1:35:65 | >0.9999 | 0.2341 | 14 |
| | | Day 1:20:80 vs. Day 7:20:80 | <0.0001 | 12.53 | 14 |
| | | Day 1:20:80 vs. Day 7:35:65 | <0.0001 | 10.06 | 14 |
| | | Day 1:20:80 vs. Day 14:20:80 | <0.0001 | 10.46 | 14 |
| | | Day 1:20:80 vs. Day 14:35:65 | 0.0002 | 9.159 | 14 |
| | | Day 1:35:65 vs. Day 7:20:80 | <0.0001 | 12.75 | 14 |
| | | Day 1:35:65 vs. Day 7:35:65 | <0.0001 | 10.27 | 14 |
| | | Day 1:35:65 vs. Day 14:20:80 | <0.0001 | 10.68 | 14 |
| | | Day 1:35:65 vs. Day 14:35:65 | 0.0001 | 9.376 | 14 |
| | | Day 7:20:80 vs. Day 7:35:65 | 0.59 | 2.315 | 14 |
| | | Day 7:20:80 vs. Day 14:20:80 | 0.7436 | 1.936 | 14 |
| | | Day 7:20:80 vs. Day 14:35:65 | 0.2839 | 3.154 | 14 |
| | | Day 7:35:65 vs. Day 14:20:80 | 0.9998 | 0.3783 | 14 |
| | | Day 7:35:65 vs. Day 14:35:65 | 0.9898 | 0.8398 | 14 |
| | | Day 14:20:80 vs. Day 14:35:65 | 0.9498 | 1.218 | 14 |
| a: ACAN | Two-way ANOVA Tukey Multiple Comparisons Test | Day 1:20:80 vs. Day 1:35:65 | >0.9999 | 0.01244 | 14 |
| | | Day 1:20:80 vs. Day 7:20:80 | 0.0332 | 4.958 | 14 |
| | | Day 1:20:80 vs. Day 7:35:65 | 0.1035 | 4.053 | 14 |
| | | Day 1:20:80 vs. Day 14:20:80 | <0.0001 | 14.52 | 14 |
| | | Day 1:20:80 vs. Day 14:35:65 | <0.0001 | 15.64 | 14 |
| | | Day 1:35:65 vs. Day 7:20:80 | 0.0327 | 4.969 | 14 |
| | | Day 1:35:65 vs. Day 7:35:65 | 0.102 | 4.065 | 14 |
| | | Day 1:35:65 vs. Day 14:20:80 | <0.0001 | 14.53 | 14 |
| | | Day 1:35:65 vs. Day 14:35:65 | <0.0001 | 15.65 | 14 |
| | | Day 7:20:80 vs. Day 7:35:65 | 0.9894 | 0.8464 | 14 |
| | | Day 7:20:80 vs. Day 14:20:80 | 0.0002 | 8.94 | 14 |
| | | Day 7:20:80 vs. Day 14:35:65 | <0.0001 | 9.995 | 14 |
| | | Day 7:35:65 vs. Day 14:20:80 | <0.0001 | 9.786 | 14 |
| | | Day 7:35:65 vs. Day 14:35:65 | <0.0001 | 10.84 | 14 |
| | | Day 14:20:80 vs. Day 14:35:65 | 0.9723 | 1.055 | 14 |
| a: COL2A1 | Two-way ANOVA Tukey Multiple Comparisons Test | Day 1:20:80 vs. Day 1:35:65 | >0.9999 | 0.001673 | 16 |
| | | Day 1:20:80 vs. Day 7:20:80 | 0.9979 | 0.5993 | 16 |
| | | Day 1:20:80 vs. Day 7:35:65 | 0.9934 | 0.7652 | 16 |
| | | Day 1:20:80 vs. Day 14:20:80 | 0.034 | 4.842 | 16 |
| | | Day 1:20:80 vs. Day 14:35:65 | <0.0001 | 13.76 | 16 |
| | | Day 1:35:65 vs. Day 7:20:80 | 0.9979 | 0.6009 | 16 |
| | | Day 1:35:65 vs. Day 7:35:65 | 0.9934 | 0.7669 | 16 |
| | | Day 1:35:65 vs. Day 14:20:80 | 0.0339 | 4.844 | 16 |
| | | Day 1:35:65 vs. Day 14:35:65 | <0.0001 | 13.76 | 16 |
| | | Day 7:20:80 vs. Day 7:35:65 | >0.9999 | 0.1659 | 16 |
| | | Day 7:20:80 vs. Day 14:20:80 | 0.0714 | 4.288 | 16 |
| | | Day 7:20:80 vs. Day 14:35:65 | <0.0001 | 13.2 | 16 |
| | | Day 7:35:65 vs. Day 14:20:80 | 0.0871 | 4.134 | 16 |
| | | Day 7:35:65 vs. Day 14:35:65 | <0.0001 | 13.05 | 16 |
| | | Day 14:20:80 vs. Day 14:35:65 | 0.0003 | 8.338 | 16 |
| a: COL1A1 | Two-way ANOVA Tukey Multiple Comparisons Test | Day 1:20:80 vs. Day 1:35:65 | 0.9914 | 0.8084 | 14 |
| | | Day 1:20:80 vs. Day 7:20:80 | 0.9871 | 0.885 | 14 |
| | | Day 1:20:80 vs. Day 7:35:65 | <0.0001 | 20.8 | 14 |
| | | Day 1:20:80 vs. Day 14:20:80 | <0.0001 | 17.72 | 14 |
| | | Day 1:20:80 vs. Day 14:35:65 | <0.0001 | 19.77 | 14 |
| | | Day 1:35:65 vs. Day 7:20:80 | >0.9999 | 0.1365 | 14 |
| | | Day 1:35:65 vs. Day 7:35:65 | <0.0001 | 21.55 | 14 |
| | | Day 1:35:65 vs. Day 14:20:80 | <0.0001 | 18.47 | 14 |
| | | Day 1:35:65 vs. Day 14:35:65 | <0.0001 | 20.52 | 14 |
| | | Day 7:20:80 vs. Day 7:35:65 | <0.0001 | 20.28 | 14 |
| | | Day 7:20:80 vs. Day 14:20:80 | <0.0001 | 17.4 | 14 |

| | | | | | |
|---|---|---|---|---|---|
| | | Day 7:20:80 vs. Day 14:35:65 | <0.0001 | 19.33 | 14 |
| | | Day 7:35:65 vs. Day 14:20:80 | 0.3709 | 2.88 | 14 |
| | | Day 7:35:65 vs. Day 14:35:65 | 0.9816 | 0.9586 | 14 |
| | | Day 14:20:80 vs. Day 14:35:65 | 0.7493 | 1.922 | 14 |
| a: COL10A1 | Two-way ANOVA Tukey Multiple Comparisons Test | Day 1:20:80 vs. Day 1:35:65 | 0.998 | 0.5891 | 14 |
| | | Day 1:20:80 vs. Day 7:20:80 | 0.2963 | 3.112 | 14 |
| | | Day 1:20:80 vs. Day 7:35:65 | 0.1646 | 3.658 | 14 |
| | | Day 1:20:80 vs. Day 14:20:80 | 0.0002 | 9.251 | 14 |
| | | Day 1:20:80 vs. Day 14:35:65 | 0.0061 | 6.261 | 14 |
| | | Day 1:35:65 vs. Day 7:20:80 | 0.1647 | 3.657 | 14 |
| | | Day 1:35:65 vs. Day 7:35:65 | 0.0861 | 4.204 | 14 |
| | | Day 1:35:65 vs. Day 14:20:80 | <0.0001 | 9.796 | 14 |
| | | Day 1:35:65 vs. Day 14:35:65 | 0.003 | 6.807 | 14 |
| | | Day 7:20:80 vs. Day 7:35:65 | 0.999 | 0.5109 | 14 |
| | | Day 7:20:80 vs. Day 14:20:80 | 0.012 | 5.742 | 14 |
| | | Day 7:20:80 vs. Day 14:35:65 | 0.3486 | 2.946 | 14 |
| | | Day 7:35:65 vs. Day 14:20:80 | 0.0233 | 5.232 | 14 |
| | | Day 7:35:65 vs. Day 14:35:65 | 0.5404 | 2.435 | 14 |
| | | Day 14:20:80 vs. Day 14:35:65 | 0.4004 | 2.796 | 14 |
| 9 a i. dsDNA | Two-way ANOVA Tukey Multiple Comparisons Test | Day 1:20:80 vs. Day 1:35:65 | 0.407 | 2.703 | 52 |
| | | Day 1:20:80 vs. Day 28:20:80 | >0.9999 | 0.04027 | 52 |
| | | Day 1:20:80 vs. Day 28:35:65 | 0.7092 | 2.022 | 52 |
| | | Day 1:20:80 vs. Day 56:20:80 | 0.8475 | 1.659 | 52 |
| | | Day 1:20:80 vs. Day 56:35:65 | >0.9999 | 0.1503 | 52 |
| | | Day 1:35:65 vs. Day 28:20:80 | 0.3946 | 2.733 | 52 |
| | | Day 1:35:65 vs. Day 28:35:65 | 0.9966 | 0.6816 | 52 |
| | | Day 1:35:65 vs. Day 56:20:80 | 0.032 | 4.432 | 52 |
| | | Day 1:35:65 vs. Day 56:35:65 | 0.2978 | 2.985 | 52 |
| | | Day 28:20:80 vs. Day 28:35:65 | 0.7039 | 2.034 | 52 |
| | | Day 28:20:80 vs. Day 56:20:80 | 0.8179 | 1.746 | 52 |
| | | Day 28:20:80 vs. Day 56:35:65 | >0.9999 | 0.197 | 52 |
| | | Day 28:35:65 vs. Day 56:20:80 | 0.1059 | 3.733 | 52 |
| | | Day 28:35:65 vs. Day 56:35:65 | 0.5987 | 2.271 | 52 |
| | | Day 56:20:80 vs. Day 56:35:65 | 0.8691 | 1.59 | 52 |
| a ii. sGAG | Two-way ANOVA Tukey Multiple Comparisons Test | Day 1:20:80 vs. Day 1:35:65 | 0.9661 | 1.132 | 52 |
| | | Day 1:20:80 vs. Day 28:20:80 | 0.0002 | 6.863 | 52 |
| | | Day 1:20:80 vs. Day 28:35:65 | <0.0001 | 9.506 | 52 |
| | | Day 1:20:80 vs. Day 56:20:80 | <0.0001 | 11.34 | 52 |
| | | Day 1:20:80 vs. Day 56:35:65 | <0.0001 | 16.07 | 52 |
| | | Day 1:35:65 vs. Day 28:20:80 | <0.0001 | 8.025 | 52 |
| | | Day 1:35:65 vs. Day 28:35:65 | <0.0001 | 10.64 | 52 |
| | | Day 1:35:65 vs. Day 56:20:80 | <0.0001 | 12.5 | 52 |
| | | Day 1:35:65 vs. Day 56:35:65 | <0.0001 | 17.26 | 52 |
| | | Day 28:20:80 vs. Day 28:35:65 | 0.3329 | 2.889 | 52 |
| | | Day 28:20:80 vs. Day 56:20:80 | 0.0232 | 4.602 | 52 |
| | | Day 28:20:80 vs. Day 56:35:65 | <0.0001 | 9.315 | 52 |
| | | Day 28:35:65 vs. Day 56:20:80 | 0.8691 | 1.59 | 52 |
| | | Day 28:35:65 vs. Day 56:35:65 | 0.001 | 6.102 | 52 |
| | | Day 56:20:80 vs. Day 56:35:65 | 0.0231 | 4.605 | 52 |
| a iii. Collagen | Two-way ANOVA Tukey Multiple Comparisons Test | Day 1:20:80 vs. Day 1:35:65 | 0.3961 | 2.772 | 21 |
| | | Day 1:20:80 vs. Day 28:20:80 | 0.0004 | 7.517 | 21 |
| | | Day 1:20:80 vs. Day 28:35:65 | <0.0001 | 8.627 | 21 |
| | | Day 1:20:80 vs. Day 56:20:80 | <0.0001 | 12.47 | 21 |
| | | Day 1:20:80 vs. Day 56:35:65 | <0.0001 | 15.18 | 21 |
| | | Day 1:35:65 vs. Day 28:20:80 | 0.0389 | 4.594 | 21 |
| | | Day 1:35:65 vs. Day 28:35:65 | 0.0054 | 5.855 | 21 |
| | | Day 1:35:65 vs. Day 56:20:80 | <0.0001 | 9.549 | 21 |
| | | Day 1:35:65 vs. Day 56:35:65 | <0.0001 | 12.26 | 21 |
| | | Day 28:20:80 vs. Day 28:35:65 | 0.8698 | 1.577 | 21 |
| | | Day 28:20:80 vs. Day 56:20:80 | 0.014 | 5.256 | 21 |
| | | Day 28:20:80 vs. Day 56:35:65 | 0.0001 | 8.133 | 21 |
| | | Day 28:35:65 vs. Day 56:20:80 | 0.2049 | 3.378 | 21 |

| | | | | | |
|---|---|---|---|---|---|
| | | Day 28:35:65 vs. Day 56:35:65 | 0.0037 | 6.091 | 21 |
| | | Day 56:20:80 vs. Day 56:35:65 | 0.3572 | 2.877 | 21 |
| b. i. dsDNA | Two-way ANOVA Tukey Multiple Comparisons Test | Day 1:MSC Only vs. Day 1:co culture | >0.9999 | 0.2706 | 16 |
| | | Day 1:MSC Only vs. Day 28:MSC Only | 0.9936 | 0.7612 | 16 |
| | | Day 1:MSC Only vs. Day 28:co culture | <0.0001 | 9.617 | 16 |
| | | Day 1:MSC Only vs. Day 56:MSC Only | 0.9987 | 0.543 | 16 |
| | | Day 1:MSC Only vs. Day 56:co culture | 0.1128 | 3.93 | 16 |
| | | Day 1:co culture vs. Day 28:MSC Only | 0.9656 | 1.115 | 16 |
| | | Day 1:co culture vs. Day 28:co culture | <0.0001 | 10.55 | 16 |
| | | Day 1:co culture vs. Day 56:MSC Only | 0.9877 | 0.8789 | 16 |
| | | Day 1:co culture vs. Day 56:co culture | 0.0513 | 4.537 | 16 |
| | | Day 28:MSC Only vs. Day 28:co culture | <0.0001 | 9.52 | 16 |
| | | Day 28:MSC Only vs. Day 56:MSC Only | >0.9999 | 0.2357 | 16 |
| | | Day 28:MSC Only vs. Day 56:co culture | 0.2067 | 3.423 | 16 |
| | | Day 28:co culture vs. Day 56:MSC Only | <0.0001 | 9.738 | 16 |
| | | Day 28:co culture vs. Day 56:co culture | 0.0041 | 6.351 | 16 |
| | | Day 56:MSC Only vs. Day 56:co culture | 0.1572 | 3.659 | 16 |
| b. ii. sGAG | Two-way ANOVA Tukey Multiple Comparisons Test | Day 1:MSC Only vs. Day 1:co culture | 0.1825 | 3.531 | 16 |
| | | Day 1:MSC Only vs. Day 28:MSC Only | 0.001 | 7.361 | 16 |
| | | Day 1:MSC Only vs. Day 28:co culture | <0.0001 | 21.82 | 16 |
| | | Day 1:MSC Only vs. Day 56:MSC Only | <0.0001 | 11.71 | 16 |
| | | Day 1:MSC Only vs. Day 56:co culture | <0.0001 | 30.97 | 16 |
| | | Day 1:co culture vs. Day 28:MSC Only | 0.0868 | 4.137 | 16 |
| | | Day 1:co culture vs. Day 28:co culture | <0.0001 | 19.79 | 16 |
| | | Day 1:co culture vs. Day 56:MSC Only | 0.0001 | 8.839 | 16 |
| | | Day 1:co culture vs. Day 56:co culture | <0.0001 | 29.63 | 16 |
| | | Day 28:MSC Only vs. Day 28:co culture | <0.0001 | 15.96 | 16 |
| | | Day 28:MSC Only vs. Day 56:MSC Only | 0.0411 | 4.702 | 16 |
| | | Day 28:MSC Only vs. Day 56:co culture | <0.0001 | 25.5 | 16 |
| | | Day 28:co culture vs. Day 56:MSC Only | <0.0001 | 11.61 | 16 |
| | | Day 28:co culture vs. Day 56:co culture | 0.0007 | 7.643 | 16 |
| | | Day 56:MSC Only vs. Day 56:co culture | <0.0001 | 20.8 | 16 |
| b iii. Collagen | Two-way ANOVA Tukey Multiple Comparisons Test | Day 1:MSC Only vs. Day 1:co culture | 0.0579 | 4.446 | 16 |
| | | Day 1:MSC Only vs. Day 28:MSC Only | 0.2263 | 3.342 | 16 |
| | | Day 1:MSC Only vs. Day 28:co culture | <0.0001 | 9.248 | 16 |
| | | Day 1:MSC Only vs. Day 56:MSC Only | 0.0002 | 8.622 | 16 |
| | | Day 1:MSC Only vs. Day 56:co culture | <0.0001 | 15.39 | 16 |
| | | Day 1:co culture vs. Day 28:MSC Only | 0.9545 | 1.193 | 16 |
| | | Day 1:co culture vs. Day 28:co culture | 0.0149 | 5.44 | 16 |
| | | Day 1:co culture vs. Day 56:MSC Only | 0.0532 | 4.51 | 16 |
| | | Day 1:co culture vs. Day 56:co culture | <0.0001 | 11.82 | 16 |
| | | Day 28:MSC Only vs. Day 28:co culture | 0.0032 | 6.544 | 16 |
| | | Day 28:MSC Only vs. Day 56:MSC Only | 0.0103 | 5.703 | 16 |
| | | Day 28:MSC Only vs. Day 56:co culture | <0.0001 | 13.01 | 16 |
| | | Day 28:co culture vs. Day 56:MSC Only | 0.9425 | 1.264 | 16 |
| | | Day 28:co culture vs. Day 56:co culture | 0.0137 | 5.499 | 16 |
| | | Day 56:MSC Only vs. Day 56:co culture | 0.0011 | 7.305 | 16 |

**Supplemental Videos:**

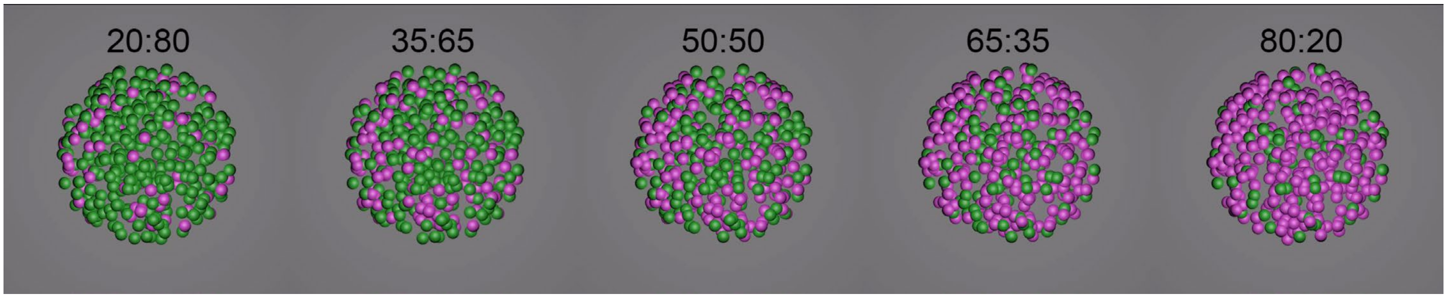


**Supplemental Video 1 | Cinema4D granular composite mixing simulations at varying microgel and spheroid ratios.**

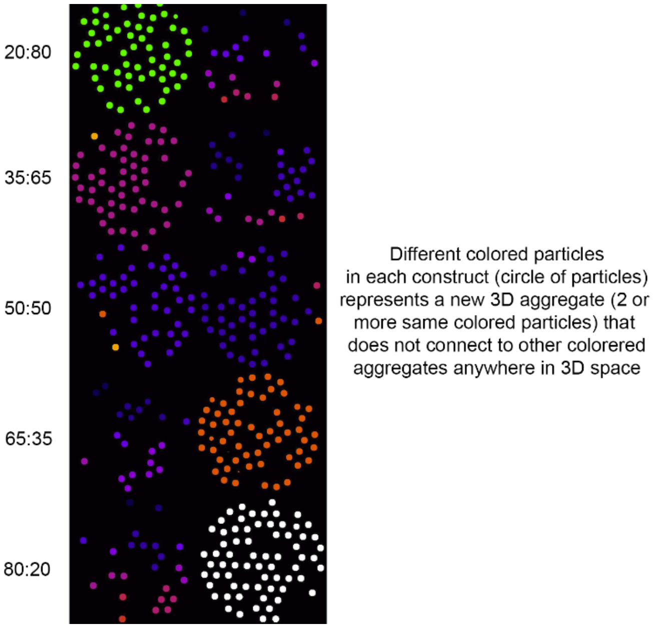


**Supplemental Video 2 | Cinema4D granular composite connectivity analysis (3d object counter).**

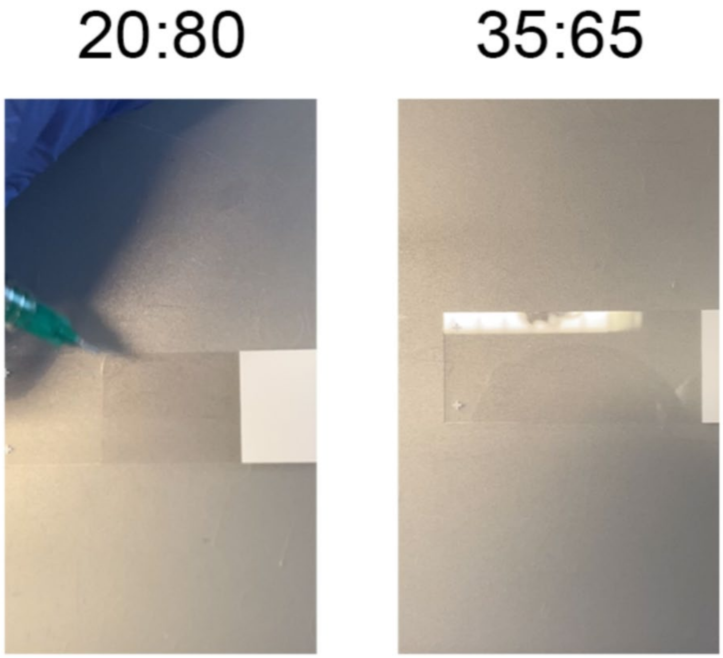


**Supplemental Video 3 | Injectability of granular composites through a 21G needle.**

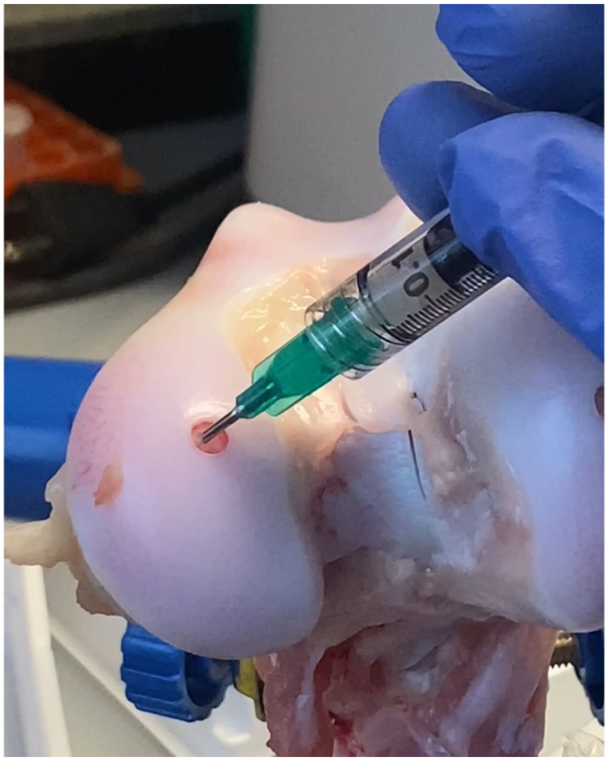

**Supplemental Video 4 | Injection of granular composite into a femoral condyle defect.**
